# Exosomes transmit retroelement RNAs to drive inflammation and immunosuppression in Ewing Sarcoma

**DOI:** 10.1101/806851

**Authors:** Valentina Evdokimova, Peter Ruzanov, Hendrik Gassmann, Syed H. Zaidi, Vanya Peltekova, Lawrence E. Heisler, John D. McPherson, Marija Orlic-Milacic, Katja Specht, Katja Steiger, Sebastian J. Schober, Uwe Thiel, Trevor D. McKee, Mark Zaidi, Christopher M. Spring, Eve Lapouble, Olivier Delattre, Stefan Burdach, Lincoln D. Stein, Poul H. Sorensen

## Abstract

Ewing sarcoma (EwS) is an aggressive childhood malignancy with a high propensity for metastasis. By analyzing cohorts of patients and age-matched healthy donors, we establish that EwS metastatic progression is accompanied by elevated plasma levels of multiple proinflammatory cytokines, interferons and extracellular vesicles (EVs). The latter were enriched with transcripts derived from LINE, SINE and ERV retroelements and from locus-specific pericentromeric regions, including HSAT2. We show that some of these RNAs, including *HSAT2* and *HERV-K*, are selectively transmitted in EwS EVs and taken up by stromal fibroblasts and peripheral blood CD33^+^ myeloid cells and CD8^+^ T-cells, inducing immune exhaustion, immunosuppressive phenotypes and proinflammatory responses. Moreover, EwS EV-derived repeat RNAs were propagated and serially transmitted in recipient cell EVs, reminiscent of viral infection. As such, this study uncovers a novel mechanism driving cancer-associated inflammation, immunosuppression and metastatic progression.

## BACKGROUND

Although a connection between inflammation and cancer was postulated by Rudolf Virchow more than a century ago^1^, it has only recently been recognized as a critical characteristic underlying tumorigenesis^2^. It is now widely accepted that cytokines, tissue remodeling enzymes, reactive oxygen and nitrogen species and other effectors of inflammation released by immune and stromal cells in the tumor microenvironment support neoplastic progression^1–3^. Emerging evidence has implicated tumor-derived extracellular vesicles (EVs), of which exosomes (∼40-150 nm vesicles) are the best studied^4, 5^, as additional prominent players in cancer-host crosstalk and cancer-associated inflammation^5, 6^. They do so by transferring tumor-derived antigens and a variety of coding and noncoding RNAs, many of which are capable of activating Toll- and RIG-like pattern recognition receptors and innate immune inflammatory responses in recipient cells^5–7^. This, in turn, induces expression and secretion of interferons (IFNs) and a variety of proinflammatory proteins, whose sustained production not only fuels tumorigenesis but also provides escape from immune surveillance, causing progressive dysfunction of the immune system^5, 8, 9^. In particular, inflammatory mediators produced by tumor and stromal cells promote generation and expansion of various immunosuppressive cells of myeloid and lymphoid origin, including myeloid-derived suppressive cells (MDSCs)^5, 10–12^, regulatory CD4^+^ (Tregs), tolerogenic and exhausted CD8^+^ T-cells^9, 13, 14^. Studying mechanisms of cancer EV-mediated inflammation and immunosuppression may therefore provide means for therapeutically reactivating key components of the immune system, restoring anticancer immunity and improving clinical efficacy of cancer immunotherapy.

Inflammation and immunosuppression may be especially relevant to Ewing sarcoma (EwS), an aggressive childhood cancer of bone and soft tissues characterized by recurrent *EWS-ETS* gene fusions, scant immune infiltrates and a high propensity for metastasis^15, 16^. It is also associated with systemic inflammation, including elevated white blood cell counts and C-reactive protein levels, each of which is correlated with poor prognosis^17^. Survival of patients with metastatic disease is poor and has not significantly improved over the past 20 years^18, 19^, in spite of high dose therapy and allogeneic stem cell transplantation^20^. Here we describe a role of EwS EVs in transmitting retroelement and pericentromeric RNAs into stromal fibroblasts and immune cells, and in expansion of immunosuppressive CD33^+^HLA-DR^-^, CD33^+^CD25^+^, CD33^+^PD-1^+^ MDSC-like and CD8^+^CD25^+^PD-1^+^ T-cells in peripheral blood of EwS patients. We also provide evidence that recipient cells become secondary transmitters of EwS-derived transcripts, by propagating and further disseminating these species in their own EVs. This phenomenon may underlie the paradoxical co-occurrence of inflammation and immunosuppression observed in EwS and, likely, in a number of other cancers.

## RESULTS

### Elevated inflammatory plasma proteins in EwS

To characterize systemic inflammation in EwS pathogenesis, we analyzed 33 plasma specimens from 30 EwS patients, 17 of which were from the Technical University München (Germany; TUM cohort) and 13 from the Institut Curie (France; EW cohort). In most cases (23 of 33), blood was drawn at diagnosis prior to chemotherapy (see Extended Data Table 1). Using the Bio-Plex profiling platform, we established that levels of 23 of 37 tested proinflammatory proteins were significantly elevated in plasma from EwS patients compared to a control cohort of 14 age-matched non-cancer subjects (Fig. 1a; Extended Data Table 2). These included tissue and bone remodeling/osteolytic enzymes such as matrix metalloproteinases (MMP-1, 2 and 3), chitinase 3-like 1 (CHI3L1) and osteopontin (OPN), secreted by tumor and certain activated immune cells, and known to play active roles in tumor invasion and inflammation. Among cytokines upregulated 2-fold or greater in EwS patients, we identified IL-2, -11, -12 [p40], -32, -35, TSLP and IL-10 family members, including IL-10, -22, -26 (Fig. 1a; p<0.0001, t-test). The upregulated cytokines are involved in host defense against pathogens, but their prolonged expression is associated with metastatic progression, cancer-associated inflammation, immune tolerance and, in the case of IL-2, IL-10 and IL-35, immunosuppression^5, 11, 12^. Also, consistent with sustained inflammation, interferon type I (IFNα and β), type II (IFNγ) and type III (IL-28A/IFNλ2 and IL-29/IFNλ1) were significantly elevated in EwS patients compared to non-cancer subjects (Fig. 1a; p<0.0001, t-test). Metastatic disease was accompanied by additional increases in OPN, CHI3L1, IL-2, IL-10 and IFNβ levels as well as other potent modulators of the immune response, such as TNFSF13, TNFRSF8, gp130 and IL-8 (Fig. 1b and Extended Data Table 1). Many of these cytokines are upregulated during persistent viral infections and cancer, affecting myeloid cells and T-cells among others^5, 8, 11^. Elevated levels of these proinflammatory proteins in patients with metastasis were independent of other clinical characteristics, such as sex, relapse or chemotherapy (Extended Data Table 3a), highlighting their strong association with EwS progression.

**Figure 1.**
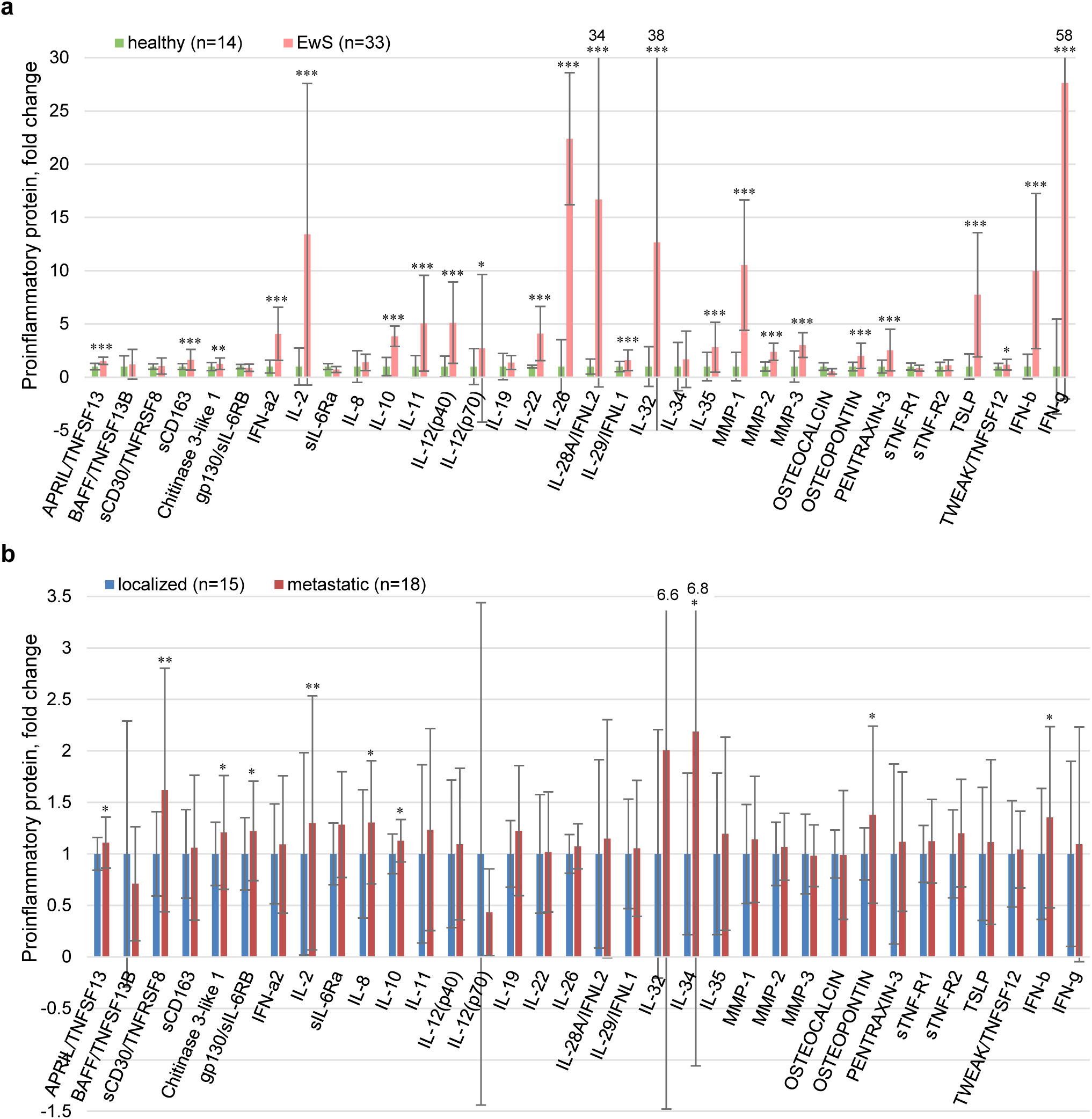
Plasma levels of proinflammatory proteins and IFNs are elevated in EwS patients. Bio-Plex 37 proinflammatory protein profiling of blood plasma from healthy donors and EwS patients, as indicated. Fold change, EwS vs healthy (**a**) or metastatic vs localized (**b**). Data are mean ± SD (n=3); *p<0.05, **p<0.005, ***p<0.0001, unpaired two-tailed unequal variance t-test.

### Predominance of repeat RNAs in EwS plasma EVs

To investigate if elevated levels of proinflammatory cytokines and IFNs might be linked to circulating EVs in EwS, plasma EVs from healthy donors and EW and TUM patients’ cohorts were purified by differential centrifugation (Extended Data Fig. 1a). Their characteristics were consistent with those of exosomes^21^, including their size (40-150 nm), enrichment with known exosome markers (CD63, CD81, Hsp70 and Hsp90), RNA-binding proteins (EWS, TIAR, hnRNPA/B, nucleolin and snRNA-associated Sm B/B’ proteins) and 20-400-nt small RNA species (Extended Data Fig. 1b-h). Moreover, plasma EV preparations were also similar to those purified from conditioned medium (CM) of EwS cell lines grown under ambient conditions or recovering after a short 30-min exposure to sodium arsenate (SA), a commonly used agent for activation of tumor-associated stress response programs, including oxidative stress^22, 23^. Notably, EVs from SA-treated cells possessed higher levels of RNA-binding proteins and Hsp70 (Extended Data Fig. 1c), likely reflective of a stress response^22^. Compared to healthy subjects (n=38), plasma EV RNA levels were significantly increased in patients with localized (n=15) and metastatic (n=19) tumors (Fig. 2a and Extended Data Tables 1, 2), being positively associated with proinflammatory protein levels in EwS patients.

**Figure 2.**
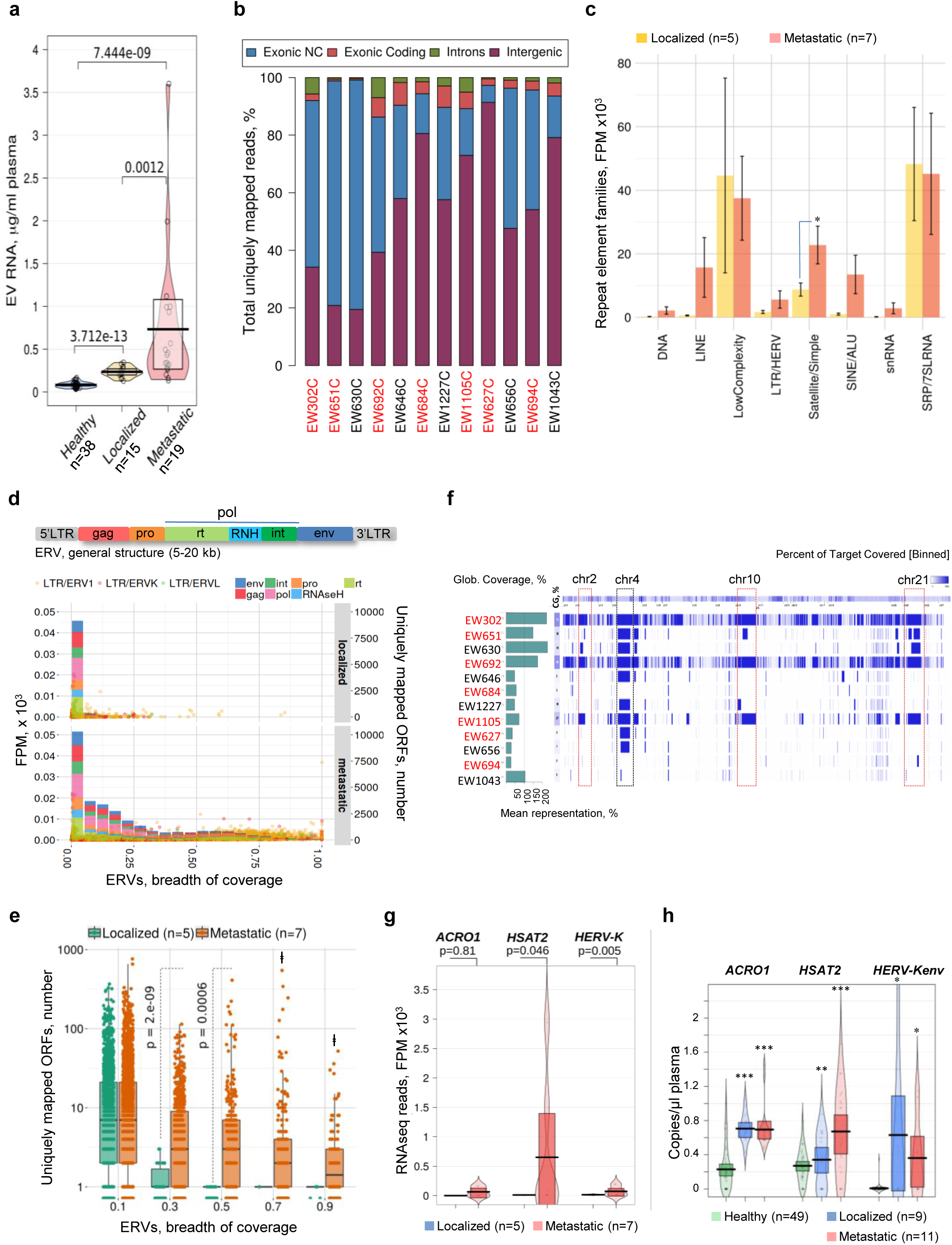
Predominance of repeat RNAs in EwS plasma EVs. **a**, Plasma EV RNA levels in healthy donors vs EwS patients, as indicated; p-values were calculated using pairwise unpaired t-test. Variance between groups was analyzed with Welch F test, p=7.0245e-14. **b,** Distribution of normalized non-ribosomal RNAseq reads in plasma EVs from 12 EwS patients; those with metastasis are marked in red. Exonic non-coding (NC), exonic, inronic and intergenic transcripts are shown. **c,** Distribution of normalized repeat RNAseq reads in EwS patients with localized vs metastatic disease, as indicated. FPM, fragments per million of uniquely mapped reads. Data are mean FPM ± SEM. *p < 0.05, Wilcoxon rank sum test. **d,** Outline of *ERV* structural organization (top) and scatter plots of ERV read coverage (bottom). The Y-axis, number of uniquely mapped ORFs; the X-axis, breadth of coverage (length covered/total length of the locus) for individual *ERV* RNAs from (**c**); see Extended Data Table 5a. Histograms, the number of unique *ERV* ORFs with different breadths of coverage. **e,** Box plots of the number of uniquely mapped *ERV* ORFs at different breadths of coverage from (**d**). Comparisons made between the number of ORFs in metastatic vs localized; p-values were calculated using Wilcoxon rank sum test; ⱡ, p-values could not be determined due to the absence of fully covered *ERV* ORFs in patients with localized disease. **f,** Genome-wide mapping of satellite repeat RNAseq reads. Blue color intensity shows breadth of coverage. Green bars to the left indicate total read counts for each sample as a percentage of mean. **g,** Quantification of selected uniquely mapped RNAseq reads for the indicated RNAs in plasma EVs from patients with metastasis vs localized tumors; p-values calculated using Wilcoxon signed-rank test. **h,** ddPCR quantification of the indicated RNAs in plasma EVs from the TUM patient cohort. Comparisons made with healthy subjects, *p<0.05, **p=0.0002, ***p<2 x 10^-8^, unpaired two-tailed unequal variance t-test. Variance between groups was analyzed with Welch F test, p=4.27e-12 (*ACRO1*), p=0.0038 (*HSAT2*) and p=0.0158 (*HERV-Kenv*).

To examine the RNA content, plasma EV RNA preparations from the EW cohort were interrogated using whole transcriptome RNA sequencing (RNAseq). Twelve of fifteen samples in this cohort yielded high-quality RNAseq libraries. Distribution of normalized uniquely mapped non-ribosomal RNAseq reads revealed a high prevalence of exonic non-coding and intergenic RNAs (Fig. 2b). Among the most prevalent species was the previously reported *7SL/SRP* RNA^24^, a core component of the signal recognition particle, as well as diverse long and short interspersed nuclear elements (*LINEs* and *SINEs*), long terminal repeat/ human endogenous retroviruses (*LTR/ERVs*) and satellite RNAs (Extended Data Table 4), many of which were elevated in patients with metastasis, although only satellite transcripts reached statistical significance (Fig. 2c and Extended Data Fig. 2a).

While most of the detected repeat-derived EV RNAs had limited or no coding capacity, some are annotated in the genome-based endogenous viral protein-coding elements (gEVE) database^25^. Based on their high coverage with respective RNAseq reads, these RNAs, including *ERV1*, *ERV2/HERV-K* and *ERV3/ERV-L* and *LINE-1* (*L1*), contain extended open reading frames (ORFs) for various retroviral polypeptides with predicted sizes ranging from 10 to more than 100 kDa, including reverse transcriptase (RT), envelop (Env), polymerase (Pol) and *L1*’s ORF1 and ORF2 (Extended Data Table 5a). Importantly, *ERVs* with extended ORFs were low or undetectable in plasma EVs from patients with localized disease, while being significantly elevated in those with metastasis (p≤0.0006, Wilcoxon test; Fig. 2d, e). Likewise, *L1* RNAs with long ORFs were significantly elevated in patients with metastasis, with the vast coming from evolutionarily younger subfamilies, including retrotransposition active human-specific L1Hs and primate L1P1, L1PA2-A5 (Extended Data, Fig. 2b-d and Table 5a). Their actual ability for retrotransposition and the capacity to produce proteins in recipient cells remains to be investigated.

Genome-wide mapping of respective RNAseq reads revealed that *L1*, *SINE*, and *LTR*/*ERV* RNAs were derived from multiple intergenic and exonic regions (Extended Data Fig. 2d, Extended Data Table 5a, b, and data not shown), potentially due to widespread epigenetic alterations of cancer genome^26^. In contrast, uniquely mapped satellite repeat reads were primarily derived from the chr2p11.2, 4p11, 10q11.21 and 21p11.2 pericentromeric regions (Fig. 2f), originating from simple (CATTC)_n_ and (GAATG)_n_ repeats, mostly located within human satellite HSAT2 and HSAT3^27, 28^. Interestingly, the majority of patients (9 of total 12) were positive for 4p11 transcripts, while RNAs mapped to all four genomic loci were found exclusively in patients with metastasis (3 of 7). These transcripts were present in both sense and antisense orientations (Extended Data Fig. 3a, b), in line with their bidirectional transcription under various stress conditions, during development, senescence and cancer^29–32^. Based on read counts, *HSAT2* and *HERV-K,* but not *ACRO1* (derived from the chr2p11.2, 4p11 and 21p11.2 regions), were significantly elevated in plasma EVs from patients with metastatic as compared to localized disease (Fig. 2g), and their increase was independent of other clinical parameters (Extended Data Table 3b).

These results were validated by droplet-based digital PCR (ddPCR), using an independent set of 20 plasma EV RNA preparations (representing 17 different TUM patients; Extended Data Table 1) and 49 age-matched non-cancer patients (Extended Data Table 2). We found that all three tested RNAs, *ACRO1*, *HSAT2* and *HERV-Kenv* (detected using primers to the relatively conserved region of the envelope gene), were significantly elevated in EwS patients compared to healthy donors (Fig. 2h). Therefore, retroelement and pericentromeric RNAs are abundant in exosome-like plasma EV particles of EwS patients, and many of them are increased with metastatic progression.

### Selective release of repeat RNAs in EVs

A complementary whole transcriptome RNAseq analysis of EVs from mock- and SA-treated A4573, TC32 and TC71 EwS cell lines revealed a profound reduction in coding RNAs (1-7% of mRNA in EVs vs greater than 20% in the respective cell lines; Extended Data Fig. 4a). By contrast, repeat RNAs, including *L1, LTR/ERV* and satellite transcripts, were elevated in EVs compared to their parental cells (Extended Data, Fig. 4b and Table 4), resembling their patterns in plasma EVs from patients with metastasis. Treatment with SA further induced levels of *LTR/ERV/ERV-K*, satellite RNAs and *7SL* RNA in their respective EVs, while *LINE*s and *SINE*s were not significantly affected (Extended Data Fig. 4b, c). Despite an increase in total levels, EVs from SA-treated cells (but not the respective parental cells) contained significantly less *ERV* RNAs with extended ORFs (Extended Data, Fig. 5c, d and Table 5b), suggesting that their release or stability in EVs are reduced under oxidative stress conditions.

Remarkably, EVs isolated from TC32 cells (but not from A4573 or TC71 cells) exhibited patterns of locus-specific pericentromeric transcripts similar to those of plasma EVs from patients with metastasis, with *(CATTC)_n_*, *(GAATG)_n_* and *HSAT2* RNAseq reads constituting up to 40% of all satellite reads in EVs from mock- and even higher from SA-treated cells, compared to less than 2% of these reads detected in respective TC32 cells (Extended Data Fig. 6a, b). Furthermore, TC32 EVs contained equal amounts of sense and antisense *HSAT* transcripts, presumably in a form of double-stranded RNAs (Jaccard coeff. ∼0.5), while only sense RNAs were detected in TC32 cells (Jaccard coeff. ∼0; Extended Data Fig. 6c), suggesting strand-specificity of their retention and release. Therefore, in contrast to *L1* and *ERV*s which were detected in all three EwS cell lines and in their EVs, *HSAT2* RNAs were expressed in only one of three EwS cell lines and preferentially released in EVs, with only a small proportion of sense transcripts retained in tumor cells, arguing for a distinct role of *HSAT2* in EwS tumorigenesis.

### EwS EVs target CD33^+^ myeloid cells and CD8^+^ T-cells

Repeat RNA-enriched plasma EVs could originate from tumor cells, as suggested by their similarity to those purified from EwS cell lines, or, alternatively, from tumor-educated platelets or immune cells^4, 6, 9^. To investigate, we employed PrimeFlow RNA assay which allows for simultaneous detection of a particular cell (based on the respective cell surface protein markers) and RNA molecules it carries. Analysis of peripheral blood mononuclear cells (PBMCs) purified from 8 healthy donors and 12 EwS patients whose plasma EVs were positive for *HSAT2* and *HERV-Kenv* (see Extended Data Table 1) revealed two *HERV-Kpol* and *HSAT2*-positive populations, CD45^+^CD33^+^ myeloid and CD45^+^CD3^+^CD8^+^ T-cells, in patient-derived PBMCs (Fig. 3a, top panels). The largest population, CD45^+^CD33^+^ cells (composed of early myeloid progenitors, monocytes and MDSCs)^10, 33^, exhibited a greater than 10-fold increase in these transcripts compared to healthy donors (p<0.0005, Wilcoxon test). Meanwhile, CD45^+^CD3^+^CD4^+^ T-cells and CD45^+^CD19^+^ B-cells were consistently negative (Fig. 3a, top panels) and their frequencies in EwS patients were reduced (Fig. 3a, bottom panels), consistent with previously reported decline of CD4^+^ T-cells in aggressive pediatric sarcomas including EwS^34^. In contrast, we observed a slight increase (∼1.3-1.5 fold) in both CD33^+^ and CD8^+^ populations (Fig. 3a, bottom panels), indicative of their potential expansion or induction in EwS patients.

**Figure 3.**
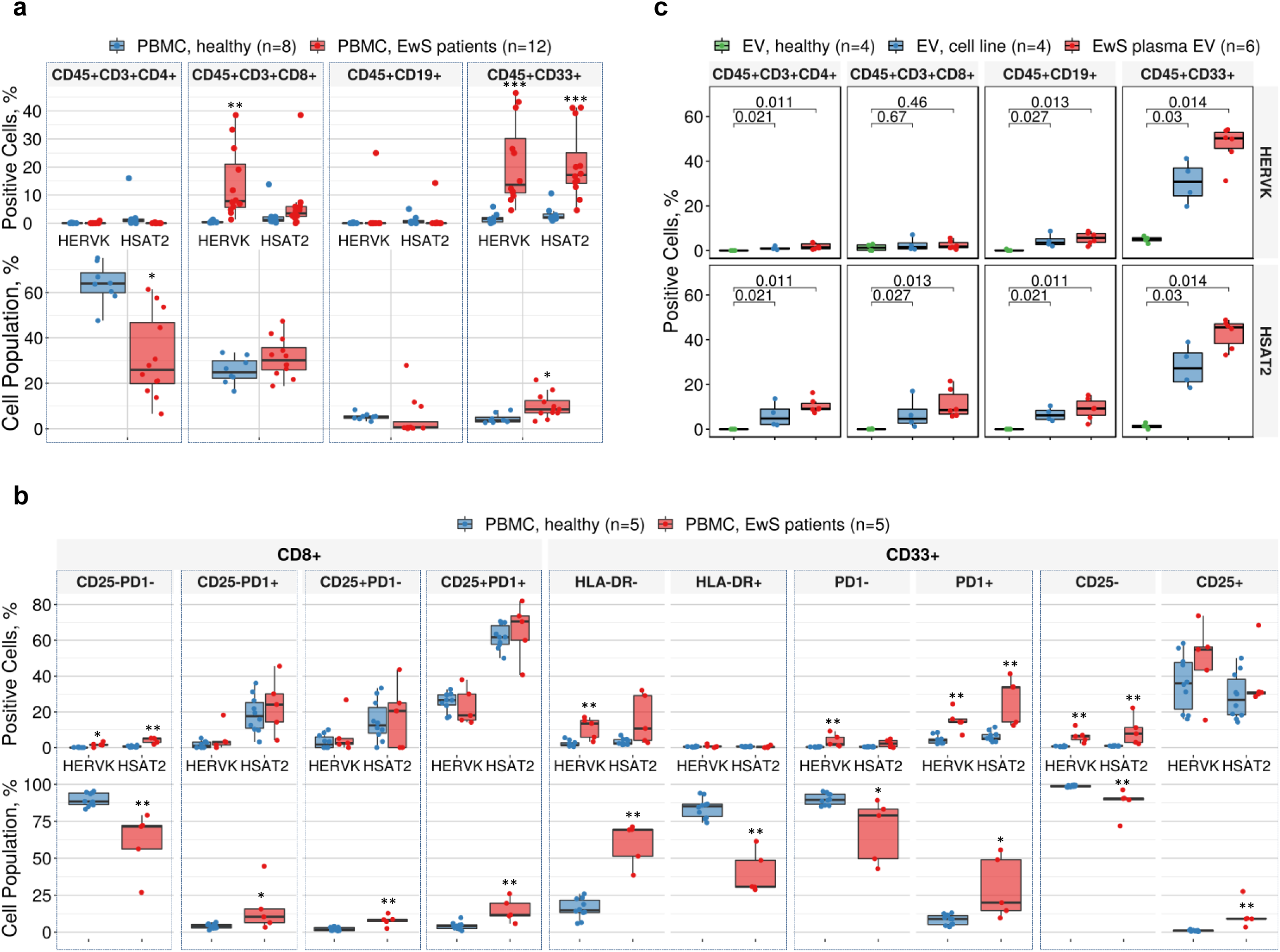
EwS EVs target immature myeloid cells and CD8^+^ T cells. **a**, Top panels, percentage of *HERV-K* and *HSAT2*-positive cells in the indicated PBMC populations from healthy donors (n=8) and EwS patients (n=12), as determined by PrimeFlow RNA assay. Bottom panels, percentage of the respective cell populations within total PBMCs. **b**, Top panels, percentage of *HERV-K* and *HSAT2*-positive cells in the indicated CD8^+^ and CD33^+^ PBMC subsets from healthy donors (n=5) and EwS patients (n=5). Bottom panels, Percentage of the respective subsets within total CD8^+^ T-cell population and within the indicated (boxed) CD33^+^ myeloid cell populations. Comparisons in (**a**) and (**b**) made with healthy donor PBMCs; *p<0.05, **p<0.005, ***p<0.0005, Wilcoxon rank sum test. **c**, Percentage of *HERV-K* and *HSAT2*-positive cells in the indicated populations of healthy donor PBMCs after 48-h treatment with EVs from plasma of healthy donors (n=4), EwS cell lines (A4573, TC32, TC71 and A673), or from EwS patients (n=6). Comparisons made with healthy donor EV-treated PBMCs; *p<0.05, **p<0.005, ***p<0.0005, Wilcoxon rank sum test.

To further characterize *HERV-Kpol* and *HSAT2*-positive populations, we employed HLA-DR (differentiation marker, to distinguish between more differentiated HLA-DR^+^ myeloid cells and HLA-DR^-^ MDSCs)^11, 33^, CD25 (CD8^+^ T-cell activation marker; also associated with MDSCs or with immunosuppressive regulatory CD4^+^ and CD8^+^ T-cells)^12, 14^ and PD-1, one of the best characterized markers of tolerance, immunosuppression and CD8^+^ T-cell exhaustion^13, 14^. Analysis of PBMCs from an additional 5 patients (all with localized tumors; pre-treatment) and 5 healthy donors identified the largest subsets positive for both *HERV-Kpol* and *HSAT2*, including CD33^+^HLA-DR^-^ (early-stage MDSCs^33^), CD33^+^PD-1^+^ and CD33^+^CD25^+^ MDSC-like cells, and CD8^+^CD25^+^PD-1^+^ T-cells with exhaustion-associated phenotype^13, 14, 35^ (Fig. 3b, top panels). In the majority of CD33^+^ subsets, levels of both *HERV-Kpol* and *HSAT2* were significantly increased in EwS patients compared to healthy donors (p<0.005, Wilcoxon test), indicating their causative association with EwS. Surprisingly however, the CD8^+^ subsets, such as *HSAT2-*positive CD25^-^PD-1^+^ and CD25^+^PD-1^-^ as well as CD25^+^PD-1^+^ (positive for both *HSAT2* and *HERV-Kpol*), were also identified in healthy donors (Fig. 3b, top panels), likely reflecting their involvement in surveillance or tolerance to self-antigens^35^. These subsets, however, were significantly expanded in EwS patients, representing only ∼1-2% of total CD8^+^ T-cell population in healthy donors and up to 15% in patients (p<0.005, Wilcoxon test; Fig. 3b, bottom panels). Likewise, the CD33^+^ *HSAT2* and *HERV-Kpol*-positive subsets were increased by ∼3-5 fold in EwS patients, while their counterpart populations were decreased (p<0.005, Wilcoxon test; Fig. 3b, bottom panels), indicating that expansion of these MDSC-like subsets could be due to altered differentiation of immature CD33^+^ myeloid cells.

To test if *HERV-K* and *HSAT2* positivity was the result of EwS EV uptake, healthy donor PBMCs were treated for 48 h with EVs from (*i*) A4573, A673, TC32 or TC71 cell lines, or from plasma of (*ii*) healthy donors or (*iii*) EwS patients. Compared to healthy donor EVs, treatment with EVs from EwS cell lines or patients plasma increased *HERV-K* and *HSAT2* levels in CD33^+^ cells by 20- and 40-fold, respectively, (p<0.05, Wilcoxon test; Fig. 3c), revealing these cells as a major target population. Similar to patients’ blood, CD8^+^ cells were the second largest affected population; however, it remains to be established if they acquire EwS EV-derived RNAs directly or via contact with the “infected” CD33^+^ antigen presenting cells. For unclear reasons, in *in vitro* settings, transfer of *HSAT2* was more efficient than that of *HERV-K* and was also observed in CD4^+^ T-cells and CD19^+^ B-cells (Fig. 3c).

To further examine the functional relevance of these findings, we employed MUTZ-3, an early progenitor myeloid cell line capable of differentiating into dendritic cells (DCs) when exposed to GM-CSF, IL-4 and TNFα^36^. We found that treatment with TC32 EVs accelerated their differentiation, including formation of cytoplasmic protrusions and elevated expression of DC-LAMP/CD208 (Extended Data Fig. 7a). However, we also observed upregulation of immunosuppressive molecules commonly produced by MDSCs in a variety of cancers^11, 12, 37^, including indoleamine 2,3 deoxygenase (IDO1) and transforming growth factor β (TGFβ; Extended Data Fig. 7a). Moreover, treatment of MUTZ-3 or HL-60 progenitor cells with EVs from A4573, TC32 or TC71 cells induced secretion of the immunosuppressive cytokines IL-10 and IL-35, along with a multitude of other proinflammatory proteins and IFNs, many of which were also elevated in plasma of EwS patients (Extended Data Fig. 7b, c, compare to Fig 1a), and may contribute to the development of functionally impaired myeloid and T-cell subtypes^12, 14, 37^. This data implicates EwS EVs in acquisition of inflammatory and immunosuppressive phenotypes by immature CD33^+^ myeloid cells and CD8^+^ T-cells, in part due to the uptake of EwS-derived repeat RNAs, including *HERV-K* and *HSAT2*.

### EwS EV RNAs are transmitted into stromal cells in vivo

To investigate if EwS EV RNAs could also be transmitted into stromal cells of the tumor microenvironment, TC32 EwS cells were implanted under the renal capsules of NOD/SCID mice, as previously described^38^. Tumor xenografts exhibited uniform sheets of small round blue cells with enlarged nuclei and minimal cytoplasm (Fig. 4a, top), characteristic of EwS^15^. ViewRNA *in situ* hybridization (ViewRNA ISH) showed that both *HSAT2* and *HERV-Kpol* were co-localized and detected in nuclei of the majority of xenografted TC32 cells (Fig. 4a, bottom). Testing for *HSAT2* in combination with probes for mouse Cd45 or Fsp-1/S100a4, markers of immune cells and fibroblasts, respectively, revealed that more than 20% of tumor-infiltrating mouse *Cd45*^+^ and *Fsp-1*^+^ cells were also positive for human-specific *HSAT2* (Fig. 4b), consistent with EV-mediated transfer from tumor cells. Neither normal human kidney tissue (Extended Data Fig. 8a) nor adjacent mouse renal epithelial cells (see asterisks in Fig. 4a, b) were positive for *HSAT2*, *HERV-Kpol, Cd45* or *Fsp-1*, confirming staining specificity and identifying tumor-infiltrating *Cd45*^+^ immune cells and *Fsp-1*^+^ stromal fibroblasts as primary targets of EwS EVs in the stroma.

**Figure 4.**
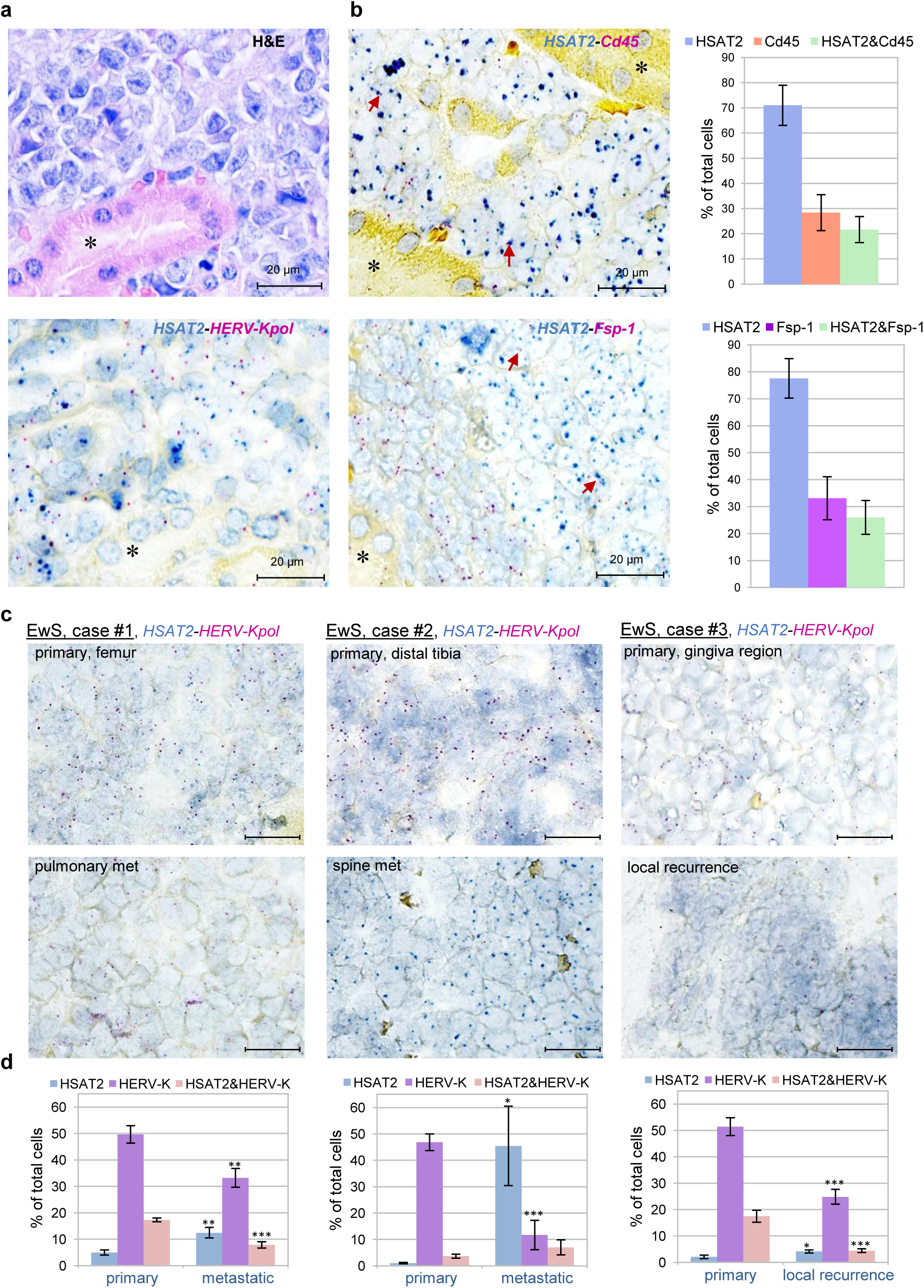
Dissemination of EwS EV RNAs in the tumor microenvironment. **a**, Haematoxylin and eosin (H&E) staining of mouse renal subcapsular TC32 EwS tumor xenografts and ViewRNA ISH staining in a corresponding serial section with the indicated probes, counterstained with hematoxylin. **b,** ViewRNA ISH with the indicated probes and associated quantification. Red arrows, co-localization of *HSAT2* (blue dots) with either *Fsp-1* or *Cd45* (purple dots) in single cells. Asterisks, adjacent normal mouse kidney parenchyma. **c, d**, Representative ViewRNA ISH images of *HSAT2* and *HERV-Kpol* expression in three matching primary and metastatic EwS tumors (**c**), and associated quantification (**d**). Data in (**b**) and (**d**) were generated using Developer XD machine learning software and expressed as means ± SD. Comparisons in (**d**) made relative to primary tumor; *p<0.05, **p<0.005, ***p<0.0005, unpaired two-tailed equal variance t-test. Scale bars, 20 μm. Additional representative images are shown in Extended Data Fig. 8.

In contrast to xenografted tumors where *HERV-Kpol* and *HSAT2* RNAs were largely co-localized, analysis of matched primary tumor and metastatic lesion biopsies from three EwS patients showed that *HERV-Kpol* and *HSAT2* RNAs were predominantly detected in localized or metastatic tumors, respectively (Fig. 4c, d and Extended Data Fig. 8b), resembling their patterns in plasma EVs (Fig. 2 h) and further implicating them into different stages of tumorigenic progression. Analysis of whole transcriptomes from 25 EwS tumor specimens (generated by Crompton et al^39^) revealed that *HERV-K* or *HSAT2* were elevated in six tumors (three of which were positive for both), and strongly associated with enrichment for an IFN-stimulated viral defense signature (p=4.15 x 10^-4^, Wilcoxon test; Extended Data Fig. 9a, b), including Toll-like receptor 7 (*TLR7*) and key IFN regulatory factors *IRF1* and *IRF7*. Therefore, repeat RNAs, including *HERV-K* and *HSAT2,* are expressed in EwS tumors and disseminated within the local tumor environment, which may contribute to activation of antiviral and proinflammatory responses in the tumor microenvironment.

### Cell-to-cell transmission of EwS EV RNAs

To further examine mechanisms driving inflammation and EwS EV RNA dissemination in the tumor microenvironment, we used MRC5 non-transformed lung fibroblasts. Treatment of these cells with TC32 EVs or mock controls for 6 h, followed by Bio-Plex 37-plex cytokine profiling of CM collected 18 h later, confirmed activation of an antiviral response, as shown by elevation of type I and type III IFNs above a 2-fold threshold (Fig. 5a). Whole transcriptome RNAseq then revealed the appearance of TC32 EV-like patterns of locus-specific pericentromeric RNA expression in recipient MRC5 cells, in contrast to mock-exposed controls (Fig. 5b). These transcripts, including *(CATTC)_n_*, *(GAATG)_n_* and *HSAT2*, were elevated by up to 10-fold in EV-treated cells compared to controls, while levels of major satellite repeats *BSR* and *MSR1 were* not changed (Extended Data Fig. 10a). In contrast to TC32 EVs in which sense and antisense transcripts were equally represented, in MRC5 cells they were found predominantly in the sense orientation (Jaccard coeff. ∼0.05; Extended Data Fig. 10b), pointing to their strand-specific accumulation or *de-novo* transcription. Among other upregulated repeat RNAs, we identified *LINE/RTE-X, LTR/ERV-L* and *SINE-Alu* (Extended Data Fig. 10c), although their functional significance remains to be determined.

**Figure 5.**
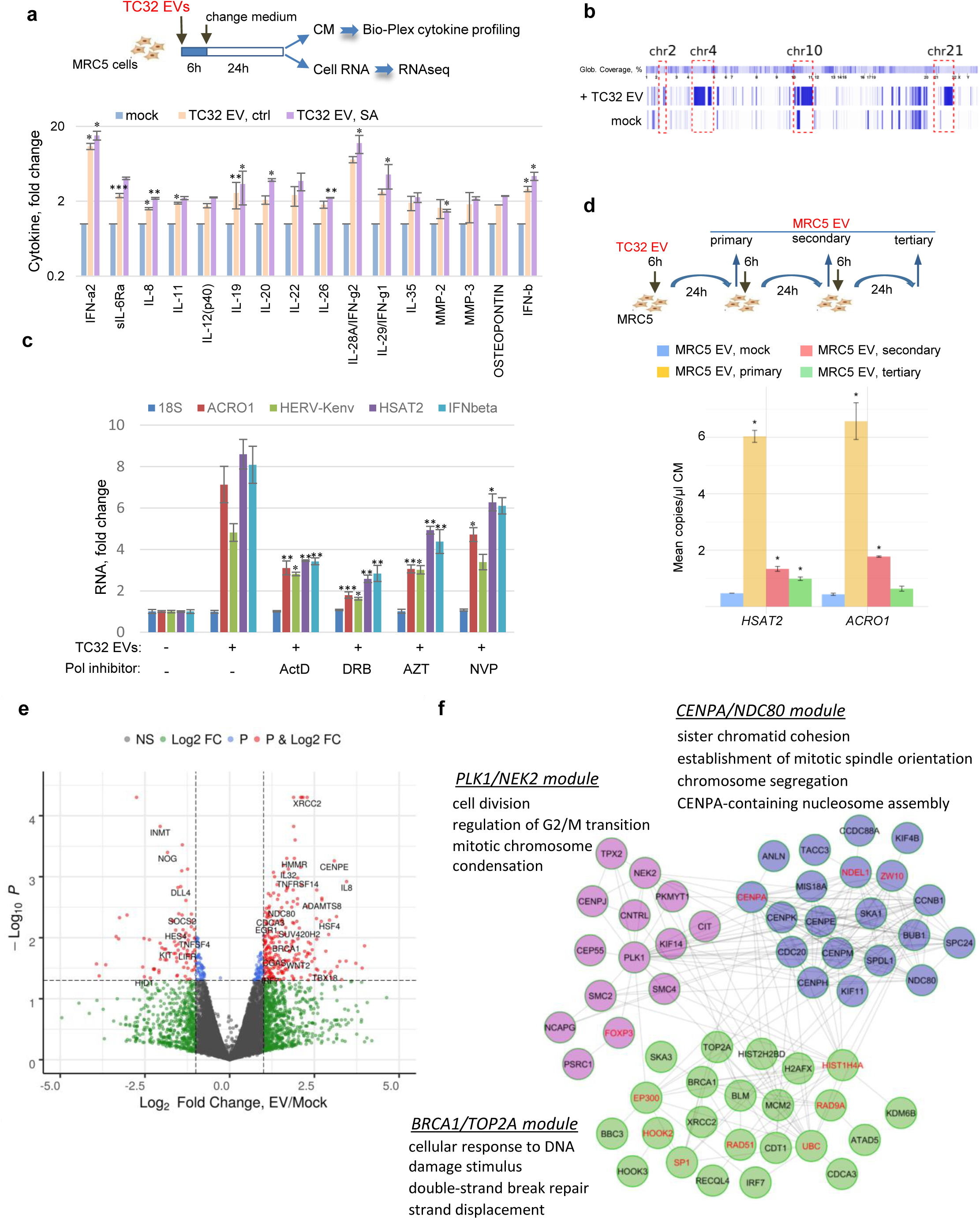
Cell-to-cell transmission of EwS EV RNAs and pericentromeric chromatin maintenance program activation in recipient cells. **a**, Outline of the experiments (top) and Bio-Plex 37 proinflammatory protein profiling (bottom), showing proteins detected in CM of TC32 EV- vs mock-treated cells above the 2-fold threshold in at least one treatment condition. Data are mean ± SD (n=3); *p<0.05, **p≤0.01; ***p<0.005, paired two-tailed t-test. **b,** Genome-wide mapping of satellite RNAseq reads from mock and TC32 EV-treated MRC5 cells. Blue color intensity shows breadth of coverage. **c**, RT-qPCR analysis of the indicated transcripts in MRC5 cells treated with TC32 EVs for 24 h in the presence or absence of Pol and RT inhibitors; 1.6 μM ActD, 200 μM DRB, 50 μM AZT, 100 μM NVP or vesicle control (-) were added for the last 8 h of 24-h treatment. Values normalized to *GAPDH* and shown as mean ± SEM (n=3). Comparisons made with TC32 EV-treated cells in the absence of inhibitors; *p<0.05, **p<0.005, unpaired two-tailed t-test. **d,** Outline of serial transfer experiments, including treatment of MRC5 cells with TC32 EVs and transfer of MRC5 EVs to freshly plated cells (top) and ddPCR quantification of the indicated EV RNAs from TC32 EV-treated cells vs mock (bottom); *p<0.05; unpaired two-tailed unequal variance t-test. Data are mean ± SEM (n=2). **e,** Volcano plot of significantly changed mRNAs in TC32 EV- vs mock-treated cells. Shown in red are mRNAs which passed p≤0.05 thresholds and showed > 2-fold expression changes. **f,** Reactome FI network analysis of mRNAs from the “Chromatin-remodeling”, “Cellular response to DNA damage” and “Cell division” GO categories upregulated in TC32 EV-treated MRC5 cells. Note that the network consists of 46 (of total 54) mRNAs and 11 externally added linkers labeled red (see full list in Extended Data Table 6c).

Strand-specific accumulation of pericentromeric transcripts in MRC5 cells prompted us to examine their potential amplification, which could be driven by Pol II^29, 31, 32^ or by RTs of either cellular (such as TERT, shown for *HSAT2*^40^) or retroviral^41, 42^ origin. We found that Pol II inhibitor DRB and, to a lesser extent, actinomycin D (ActD; a pleiotropic polymerase inhibitor) strongly blocked accumulation of *HSAT2, ACRO1* and *HERV-Kenv,* along with *IFNβ* mRNA used as an internal control (Fig. 5c). The RT inhibitors 3’-Azido-3’-deoxythymidine (AZT/zidovudine; an inhibitor of *HIV*, *HERV-K* and *L1*-encoded RT activities^42, 43^) and nevirapine (NVP) had moderate effects, with AZT being more efficient in reducing levels of *HSAT2, HERV-Kenv* and *ACRO1* by ∼1.6 and 2.5-fold, respectively (Fig. 5c); the observed reduction in *IFNβ* mRNA levels could be due to a decrease in repeat RNAs and diminished antiviral response under these conditions. Neither treatment affected levels of 18S rRNA, indicating specificity and supporting the involvement of Pol II- and, potentially, RT-dependent mechanisms into propagation of EwS EV-derived RNAs in recipient cells. Furthermore, recipient MRC5 cells were also capable of releasing *HSAT2* and *ACRO1* in their own EVs and transferring them to freshly plated MRC5 cells. Strikingly, even after a third transfer, *HSAT2* levels were significantly higher than those in EVs from mock-treated MRC5 cells (p<0.05; t-test; Fig. 5d), indicating that “infected” fibroblasts are capable of amplifying and further disseminating EwS-derived transcripts.

Further analysis of RNAseq data revealed that in addition to repeat RNAs, there were small but intriguing changes in coding transcripts. Among 109 mRNAs downregulated in TC32 EV-treated MRC5 cells relative to mock, we identified suppressor of cytokine signaling 2 (*SOCS2*), transcriptional regulators *HES4* and *KLF2*, proto-oncogene receptor tyrosine kinase *KIT* and the Notch ligand *DLL4* (Fig. 5e and Extended Data Table 6a). Among 297 upregulated mRNAs, there were *IRF7, IL8, IL32, TNFRSF14* and *MB21D1/CGAS* (Fig.5e and Extended Data Table 6b), consistent with the observed activation of antiviral and proinflammatory responses. However, the largest group of upregulated mRNAs (54 of 297) were those encoding centromeric proteins and kinetochore complex assembly components (“CENPA/NDC80” module), key regulators of centrosome maturation, spindle assembly and chromatin condensation (“PLK1/NEK2” module), and other cell division and DNA damage control proteins (“BRCA1/TOP2A” module; Fig.5f and Extended Data Table 6c), all of which are known to be involved in pericentromeric heterochromatin maintenance^29, 44, 45^. Activation of these pathways may thus be indicative of extensive centromeric heterochromatin remodelling in recipient cells. Collectively, our findings demonstrate transfer of pericentromeric and retroelement RNAs in EwS EVs, their propagation in recipient cells, and further transmission in the recipient cell EVs.

## DISCUSSION

Our study shows that global de-repression of constitutive heterochromatin regions, a process that has previously been linked to different stages of development, aging and cancer^26, 29, 31, 32^, may have more far-reaching implications than previously anticipated. In particular, we establish that diverse classes of retroelement and pericentromeric chromatin-derived transcripts including *L1*, *HERV-K, HSAT2* and *ACRO1* are not only massively expressed in EwS cell lines and tumors, but are also preferably secreted in EVs and their levels in plasma EVs are associated with inflammation and metastatic progression. Our data provides strong support for the Trojan exosome hypothesis^46^, which postulates that HIV and other retroviruses and LTR (and non-LTR, as shown here) retrotransposons exploit exosomal pathways for their spread, preferentially infecting cells of the immune system. Indeed, we show that EwS EV-derived RNAs are taken up by healthy donor PBMCs, predominantly targeting CD33^+^ myeloid cells and, to a lesser extent, CD8^+^ T-cells. Respective *HERV-K* and *HSAT2-*positive cell populations were significantly expanded in the peripheral blood of EwS patients compared to healthy donors. These populations exhibited phenotypic features of MDSCs (CD33^+^HLA-DR^-^, CD33^+^CD25^+^, CD33^+^PD1^+^)^11, 33, 37^ and of exhausted or regulatory CD8^+^ T-cells (CD8^+^CD25^+^PD-1^+^)^13, 35^. EwS-derived RNAs may thus co-opt key cell populations of the innate and adaptive immune system, skewing them towards immunosuppressive phenotypes. This phenomenon may underlie the reported EV-driven generation of MDSCs^6, 9, 11^, and their expansion in blood of patients with aggressive sarcomas including EwS^34^, melanoma and other types of cancer^11^. Induction of these immunosuppressive cells and their ability to produce central mediators of inflammation and tolerance (including IL-10, IL-35, IDO1 and TGFβ detected in patients’ blood and in *in vitro* models used in this study) may further compromise proper differentiation of myeloid cells and lymphocytes, subsequently dampening antitumor immunity.

Improperly differentiated immune cells may also infiltrate or be generated at tumor sites, which could contribute to the observed scarcity of mature DCs and cytotoxic CD8^+^ T-cells in EwS tumors^16, 17^. These findings may have implications for a wider range of immunologically cold tumors, given the reported association between high expression of *L1*, *ERVs* and *HSAT2* and immunosuppression, characterized in part by fewer infiltrating CD8^+^ T-cells in *HSAT2-*high colon and pancreatic cancers^47^.

Given the multiple virus-like features of *LINEs*, *SINEs*, *LTR/ERVs* and other repeat-containing RNAs, such as exposed 5’-triphoshate ends^7^, EV RNA unshielding^24^, pathogen-like motif patterns *(i.e.,* high CpG content in AU-rich contexts as shown for *HSAT2*^48^), as well as their ability to generate dsRNAs and/or RNA-DNA intermediates^40, 41, 49–51^, it is not unexpected that we observed vigorous IFN-driven antiviral responses triggered by EwS EVs in the recipient cells, including myeloid progenitors and fibroblasts. Because of the high abundance of hundreds of thousands of such species detected in EwS EVs in this study and in other cancers^7, 52^, this may be a ubiquitous process which is not restricted to particular retroelement species. Most likely, these species act as a broad population with a common action, each of which is capable of triggering innate responses in a wide range of host cells. Their synergistic effect may create a permissive tumor environment both locally and systemically. This does not exclude, however, that some of them, including *HERV-K* and *HSAT2*, may have more specialized functions, such as selective targeting of immature myeloid cells and CD8^+^ T-cells to induce immunosuppressive phenotypes.

Another unexpected finding of this study is that despite their obvious lack of coding or replicative potential, some of these transcripts, including *HSAT2* and *ACRO1*, are further propagated in recipient MRC5 fibroblasts and then released in their respective EVs, being capable of “infecting” freshly plated cells. They seemingly exploit cellular machinery of recipient cells for their own propagation, packaging and further disseminating in recipient cell EVs. Although the exact mechanism remains to be determined, our data indicates that it can be mediated by Pol II, based on the efficient blockade of retroelement RNA accumulation in recipient cells by the RNA Pol II inhibitor DRB. One possibility is that these RNAs recognize and remodel the respective heterochromatin regions to activate transcription. This is supported by the observed upregulation of mRNAs whose protein products are responsible for pericentromeric chromatin maintenance as well as by multiple reports showing that pericentromeric chromatin transcription is Pol II-dependent and regulated by pericentromeric transcripts and by their ability to recruit chromatin remodelers^29, 31, 32^. Alternatively, retroelement activation in recipient cells could be triggered by antiviral and stress responses known to induce transcription of endogenous retroelements, including *HERV-K*, *L1* and pericentromeric transcripts^29, 31, 32, 41^. Also, we cannot exclude reverse transcription as a potential mechanism, considering that some *L1, HERV-K* and other *LTR/ERVs* identified in EwS EVs may encode RT and other retroviral proteins, and because RT inhibitors were moderately effective in blocking accumulation of these transcripts in MRC5 cells. None of these scenarios are mutually exclusive, nor require fully intact *L1* or *HERV-K,* or their reintegration into the genome, as would be expected from canonical models^26, 41, 49, 53^. Further investigation into these mechanisms may provide unique insights into cancer-associated inflammation, evasion of antitumor immunity and immunosuppression, and open novel directions for therapeutic targeting.

## METHODS

### Plasma EV purification and analysis

Blood plasma specimens were obtained from Ewing sarcoma participants enrolled in GenEwing and NEO-IDENT studies, as well as from non-cancer individuals undergoing routine checkups at the allergy and asthma clinic at the Technical University München. Each cohort received approvals from the respective institutional review boards, Comité de Protection des Personnes, Ile-de-France (#09-12037) and the Ethikkommission der Medizinischen Fakultät der Technischen Universität München (#2562/09). After thawing, plasma (1 ml/patient) was clarified by centrifugation at 10,000 g for 20 min. Supernatants were passed through 0.22 μm filters, diluted with an equal volume of ice-cold phosphate buffered saline (PBS) and centrifuged at 100,000 g (S110-AT rotor, Thermo Scientific), 4°C for 120 min. Pellets were re-suspended in 3 ml of PBS and pelleted again by ultracentrifugation, to remove plasma contaminants. Purified EVs were either directly used for RNA isolation or dissolved in 300 μl PBS and used for immunoblotting (see Extended Data Table 7 for antibodies used) or Nanoparticle Tracking Analysis (NS300 NTA; NanoSight). Respective protocols for clinical sample handling and processing were reviewed and approved by the University of Toronto ethics committee (#35129).

### EV purification from cell cultures and functional assays

Cell line sources and growth conditions are listed in Extended Data Table 8. For EV purification, EwS cell lines were grown on four 15-cm plates up to ∼80% confluency in 10% fetal bovine serum (FBS) medium, treated with 0.5 mM sodium arsenite (SA) for 30 min or PBS, and then the medium was replaced with fresh one containing 2% EV-depleted FBS (System Biosciences). Conditioned medium (CM) was collected after overnight incubation and subjected to sequential centrifugation at 1,500 g for 10 min and 10,000 g for 20 min, to remove cell debris and larger particles. CM was then concentrated using the Amicon Ultra-15mL-30K tubes (Millipore), passed through 0.22 μm filters, diluted with equal volumes of PBS and subjected to ultracentrifugation at 100,000 g for 120 min. To remove contaminating proteins, EV pellets were then re-suspended in 3 ml of PBS and pelleted again by ultracentrifugation. Purified EVs were dissolved in 500 μl PBS and used for functional testing.

Treatment of MRC5 and HL-60 cells with 50 μl EVs (1x10^9^ particles/100,000 cells/well in 24-well plates) or mock isolates prepared from the equal volume of 2% FBS non-conditioned growth medium was carried out in 500 μl of the appropriate 2% FBS growth medium for 6 h, in duplicate. Cells and CM were collected after 18 or 24 h incubation in a fresh medium, as indicated in figure legends. Where indicated, polymerase (ActD, #A1410 and DRB, # D1916, Sigma) or reverse transcriptase (AZT, #A2169 and NVP, #SML0097, Sigma) inhibitors were added for the last 8 h of 24-h incubation of MRC5 cells with mock or TC32 EVs. Cell-to-cell transfer was assayed in the same 24-well format (12 wells/condition), by applying EVs isolated from CM onto freshly plated MRC5 cells for 6 h and collecting CM 24 h later. Due to low EV yields from mock-treated MRC5 cells, double amounts of cells were used for their purification. After optimizing conditions, each experiment was repeated at least three times.

### Bio-Plex proinflammatory protein profiling

Proinflammatory protein profiling of blood plasma (12 μl/reaction) or CM from the cell lines (50 μl/reaction) was done using the magnetic bead-based multiplex immunoassay Bio-Plex 200 system and the 37-plex human inflammation panel (171AL001M, Bio-Rad) following manufacturer’s recommendations. Each CM sample was prepared in duplicate and experiments were repeated at least three times using different EV preparations. Plasma proinflammatory protein profiles were generated from three independent measurements of the same specimens. Standard and sample cytokine concentrations were then calculated using the Bio-Rad Bio-Plex 200 software. Background corrected signals for each cytokine were normalized to the mean signal intensity obtained for each sample and were plotted using the mean and standard deviation from three independent experiments.

### MUTZ-3 differentiation and EV treatment

Immature and mature MUTZ-3 DCs were generated as described^36^. Briefly, MUTZ-3 were cultured in MEMα growth medium (12571-063, ThermoFisher) containing 20% FBS and 40 ng/ml GM-CSF (PHC2015, ThermoFisher). To generate immature DCs (iDCs), MUTZ-3 cells were incubated for 7 days in a 6-well plate (1.5x10^6^ cells/well) in 2 ml of growth medium supplemented with 100 ng/ml GM-CSF, 20 ng/ml IL-4 (P05112, R&D Systems) and 2.5 ng/ml TNFα (P01375, R&D Systems) in the presence or absence of TC32 EVs (100 µl/2x10^9^ particles/well) for the first 3 days of incubation. Maturation was induced by adding a cytokine cocktail consisting of 50 ng/ml TNFα, 100 ng/ml IL-6 (Q75MH2, R&D Systems), 25 ng/ml IL-1β (NP000567, R&D Systems) for the additional 3 days.

### PBMCs treatment, PrimeFlow RNA assay and flow cytometry

Bio-banked PBMCs (∼2-4×10^6^ cells) purified by Ficoll density gradient centrifugation from EDTA tube-collected whole blood of EwS patients and healthy individuals under the Ethikkommission der Medizinischen Fakultät der Technischen Universität München protocol (#2562/09) were either assayed directly, or pre-treated with EVs from plasma of healthy donors, EwS patients or EwS cell lines. Treatment of healthy donor PBMCs (2x10^6^ cells/well in 12-well plates, in duplicates) with 100 μl EVs (2x10^9^ particles/well) was carried out in 1 ml of RPMI-1640, 10% FBS growth medium for 48 h, and then cells were processed for staining using the PrimeFlow RNA kit (#8818005204; ThermoFisher). Staining was done according to the manufacturer’s instructions, stepwise, using cell viability dye (FVD EF506), fluorescently-labeled antibodies against the respective cell surface markers and, finally, RNA probes for HERV-K AF647 (Type 1) and HSAT2 AF750 (Type 6). Positive control probe RPL13a AF647 (Type 1) was used for optimization; see Extended Data Table 7 for the full list of reagents used. For flow cytometry, cells were re-suspended in 200 μl of staining buffer (554657, BD Pharmingen) and analyzed using a BD Fortessa X-20 SORP (BD Biosciences) and BD FACSDiva 8.0.2 software. Healthy donor and EwS patients’ samples were run in parallel. Positive events were identified based on isotope control antibodies, fluorescence minus one (FMO) controls and excluding non-viable FVD-positive cells and cell aggregates. Gating strategy is shown in Extended Data Fig. 11.

### ViewRNA ISH

Archived formalin-fixed paraffin-embedded (FFPE) sections of EwS tumors were obtained from the Technical University of Munich School of Medicine Biobank with approval from the Institutional Review Board of the TUM School of Medicine (#2562/09). FFPE sections of mouse xenografts of TC32 EwS cells were provided by Dr. A.M. El-Naggar and obtained as described earlier^38^. ViewRNA ISH was performed using ThermoFisher reagents, including ViewRNA ISH tissue 2-Plex assay kit (QVT0012) and probe sets designed to specifically hybridize to human HERV-K*pol* (VF1-18759-06) and HSAT2 (VA6-19493-06), and mouse S100a4/Fsp-1 (VB1-18990-06) and Ptprc/Cd45 (VB1-12657-06) RNAs, in accordance with the manufacturer’s instructions. Briefly, 5 μm FFPE sections were baked and deparaffinised, followed by heat pretreatment and protease digestion. Target probes were then hybridized for 2 h at 40°C followed by ViewRNA signal amplification. Signal detection was performed sequentially, using Fast Blue and Fast Red substrates, and slides were then counterstained with hematoxylin. Signal was considered specific if normal kidney tissue and non-probe controls stained in parallel showed less than 1 dot per 20 cells. Regions of interest were captured at x100 magnification using a Leica DM2000LED microscope and LAS software with a pixel resolution of 0.04 μm/pixel. Stain separation of the fast blue, fast red and nonspecific background signals were achieved using a python implementation of the independent component analysis approach^54^. The resulting deconvoluted images were analysed using a custom rule set built in the object-oriented machine learning software Developer XD (Definiens AG, Munich, Germany). The rule set first separated nuclear regions from nonspecific signal using a weighted combination of all isolated stain intensities, with watershed and object reshaping algorithms being employed to identify and separate closely-spaced nuclei. Blue and red spots were then detected with the fast blue and fast red channels, respectively. The resulting cell objects were exported as a table listing the number and type of spots within each cell. The number of cells containing at least one red spot, one blue spot, or both were tabulated as a fraction of total cells detected.

### RNA extraction and whole transcriptome RNAseq

Total RNAs from EwS cell lines and the respective EVs, and plasma EVs were purified using the *mir*Vana RNA isolation kit (AM1561, ThermoFisher), treated with Turbo DNase (AM1907, ThermoFisher) and quantified using the Qubit RNA high sensitivity kit (Q32852, ThermoFisher). RNA integrity was analyzed with the RNA 6000 Pico or Small RNA chips (Agilent Technologies), and preparations of the sufficient quantity and quality were then used to construct strand-specific RNAseq libraries with the TruSeq Stranded Total RNA library prep and RiboZero rRNA removal kit (20020596, Illumina). Libraries were pooled in sets of 4 and sequenced using the HiSeq PE cluster kit v4 on the Illumina HiSeq 2500 sequencer.

### Reverse transcription (RT), quantitative (qPCR) and droplet-based digital PCR (ddPCR)

cDNA was synthesized using SuperScript III Reverse Transcriptase (18080044, ThermoFisher), 0.5 μg of DNase-treated RNA and random hexamer primers. Negative control RNA samples were pretreated with a mixture of RNase A and T1, to control for genomic DNA contamination. One μl cDNA (of total 20 μl) was used for PCR with the respective primer sets (Extended Data Table 9) and Q5 High-Fidelity 2X Master Mix (M0492S, NEB). PCR reactions were incubated at 94°C for 1 min followed by 4 cycles of 94°C for 30 s, 58°C for 45 s and 72°C for 20 s, and by additional 30 cycles of 94°C, 30 s, 54°C, 45 s and 72°C, 20 s, with a final incubation at 72°C for 4 min. PCR products were analyzed on 1.5% agarose gel and confirmed by Sanger sequencing.

Quantitative PCR (qPCR) was performed using 1 μl cDNA, respective primers and the PowerUp SYBR Green Master Mix (A25777, ThermoFisher) on the ViiA 7 Real-Time PCR system (ThermoFisher) according to the manufacturer’s recommendations for the standard cycling mode with primers with T_m_<60°C. A non-template control was included in all qPCR reactions. After optimizing experimental conditions, each experiment was independently repeated at least three times in duplicate.

ddPCR was performed on the QX100 Droplet Digital PCR system (Bio-Rad) using 1 μl cDNA, the respective primer sets and the QX200™ ddPCR™ EvaGreen Supermix (1864033, Bio-Rad). PCR reactions were incubated at 95°C, 5 min, followed by 40 cycles of 95°C for 30 s and 58°C for 1 min, with a final incubations at 4°C for 5 min and at 90°C for 5 min. The average number of accepted droplets of the valid measurement results was ∼17,000. Data from the droplet reader presented as copies per 1 μl PCR reaction were converted to copies per ml CM or plasma based on the known equivalents of input RNA. Visualization and standard error calculation were performed in R.

### Bioinformatics and statistical analysis

Paired-end strand-specific 100 or 125-bp reads were aligned to human genome build 37 (hg19/GRCh37) using the BWA aligner v.0.7.13 with subsequent filtering of ribosomal and tRNA reads, followed by removal of secondary alignments and reads with mapping score less than 30. Transcript expression was quantified as the numbers of fragments per million (FPM). Based on annotations from the Known Genes track of UCSC browser, reads were assigned to different transcript classes using bedtools suite 2.17.0. Analysis of repeat RNAs was done with hg19-compliant RepeatMasker track annotations. To quantify expression of different transcript classes, overlapping annotations were resolved giving priority to features in a hierarchical order: miRNA > piRNA > lincRNA > Repeatmasker annotations > UTRs > exon > intron. The annotated reads were partitioned into four bins: exonic coding, exonic non-coding (including *7SL* RNAs, snRNAs, lncRNAs, 5’ and 3’UTRs), intronic and intergenic with the results plotted using R statistical environment (https://www.R-project.org/). Coverage for each transcript was calculated with bedtools, and figures were generated using the ggplot2 package^55^. No statistical methods were used to predetermine sample size. All statistical analyses were performed using R with packages obtained from Bioconductor (https://www.bioconductor.org) and/or from CRAN (https://cran.r-project.org/web/packages/). RNAseq and ddPCR data were analyzed with Shapiro-Wilk test (to verify normal distribution of data), followed by the unpaired t-test on log-transformed values with subsequent Benjamini & Hochberg (BH) adjustment. For data that failed a normality test, the Wilcoxon rank sum test was applied. For comparisons among more than two groups of samples, one-way ANOVA or Welch F test (in cases when homoscedasticity condition was violated) were used. All custom codes used for statistical analysis and visualization are available upon request.

### Differential expression, Gene Ontology (GO) and Reactome Functional Interaction (FI) network analysis

Differential expression analysis of protein-coding genes was done by aligning RNAseq reads with STAR 2.5.1b (https://github.com/alexdobin/STAR) and running Cufflinks 2.2.1 (http://cole-trapnell-lab.github.io/cufflinks/) using Ensembl gene annotations (release 87). Transcripts showing 2-fold difference and p-value < 0.05 were considered as differentially expressed. Gene ontology analysis (GO) and functional enrichment in the molecular function, biological process and cellular component categories were performed using DAVID, Panther (http://www.pantherdb.org/) and GO (http://geneontology.org/) online databases. Highly enriched “Chromatin-remodeling”, “Cellular response to DNA damage” and “Cell division” GO categories were further analyzed using the Reactome (http://www.reactome.org) database and Reactome FIViz Cytoscape plugin^55^, to identify network modules of highly connected genes and their significantly enriched (FDR<0.05) biological processes.

### Data availability

RNAseq data generated from 12 EwS patients’ plasma EVs and from A4573, TC32 and TC71 EwS cell lines, mock- or SA-treated, and their respective EVs were deposited at the Sequence Read Archive (SRA) under the accession number PRJNA548159 (pending). Source data for the figures and reagents are provided with the paper and are also available from the corresponding authors upon request.

## Acknowledgements

We are thankful to Victoria Luchenko, a wonderful person and devoted researcher at the National Institutes of Health (NIH) who recently lost her fight against cancer, for inspiration and fruitful discussions. We also would like to thank all patients and their families, and the personnel of the Genetic Somatic Unit and the Institute of Pathology for their help in collecting blood samples. We thank B.D. Crompton and K. Stegmaier for sharing their RNAseq EwS datasets, C. Lachance and S. Wells for assistance with Bio-Plex and ddPCR experiments, M. Shure, B. Samelson, A. Patel, E. Ortenberg and M. Bourbonnière for ViewRNA ISH and PrimeFlow RNA assay support, and A.M. El-Naggar for xenografted EwS sections. The authors would like to acknowledge the Spatio-Temporal Targeting and Amplification of Radiation Response (STTARR) program and its affiliated funding agencies. This work was primarily supported by funds from CIHR Foundation grant FDN-143280 and from the British Columbia Cancer Foundation through generous donations from Team Finn and other generous riders in the Ride to Conquer Cancer (to PHS). This work was also supported by grants from Cura Placida Children’s Cancer Research Foundation (to PHS, UT, and SB), the Wilhelm-Sander Stiftung (2009.901.3, to SB), the Federal Ministry of Education and Research (BMBF) Germany (PROVABES; 01KT1311, to SB), and the Government of Ontario (to VE, PR, SHZ, VP, LEH, JDM and LDS).

## Author Contributions

VE conceived and designed the study, performed the experimental work and co-wrote the manuscript; PR conducted bioinformatics analysis, data interpretation, preparation of figures, legends and relevant sections in the Methods; HG, KS, KS, UT, SS and SB provided clinical specimens and contributed to ddPCR, PrimeFlow RNA and ViewRNA-ISH experiments; EL and OD provided human samples; SHZ helped to design, execute and interpret RT-qPCR data; VP advised on cytokine profiling experiments, immunohistochemistry and immune cell models; MOM performed Reactome pathway analysis; JDM and LH supervised RNAseq and data analysis; TDM and MSZ performed ViewRNA image spot analysis and wrote the relevant section in the Methods; CMS provided help with flow cytometry and PrimeFlow RNA data analysis; SB, LDS and PHS advised on experimental design, data analysis and interpretation and manuscript writing; PHS coordinated the project and secured funding. All authors edited and approved the manuscript.

## Author Information

The authors declare no competing financial interests. Correspondence and requests for materials should be addressed to VE and PHS.

## SUPLEMENTARY FIGURES

**Extended Data Figure 1.**
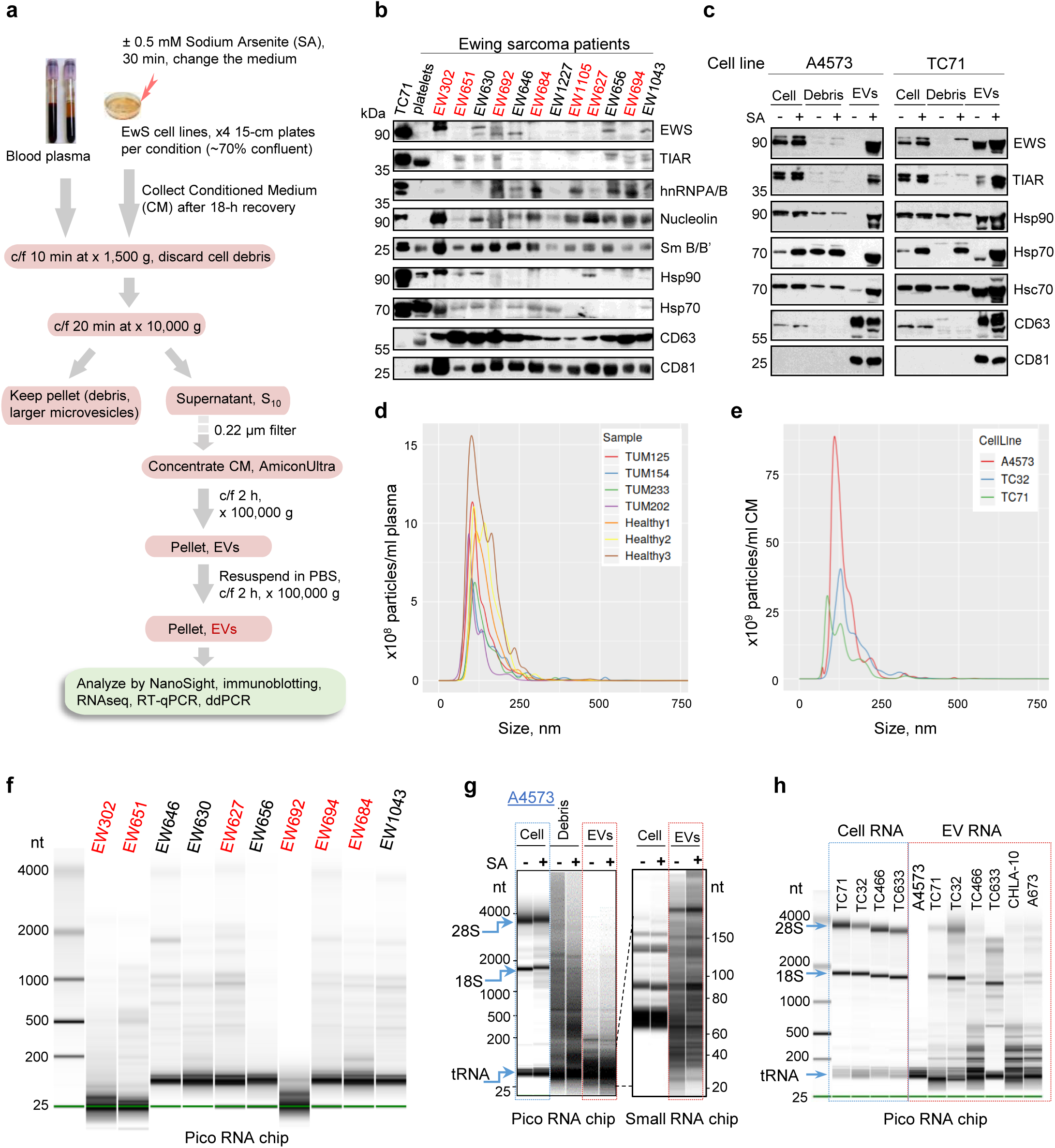
EV purification and characterization. **a**, Outline of EV purification from blood plasma and cell lines. **b, c**, Immunoblotting of EV preparations from patient blood plasma (patients with metastasis are marked red), and from EwS cell lines treated with PBS (-) or 0.5 mM sodium arsenate (SA) (**c**). Shown in (**c**) are total cell extracts (Cell), larger vesicles and cell debris after x10,000 g centrifugation (Debris) and purified EVs. **d, e,** EV size distribution by nanoparticle-tracking analysis of EVs from plasma of healthy donors and EwS (TUM) patients (**d**), and from the indicated cell lines (**e**). **f-h**, Analysis of EV RNAs purified from EwS patients’ plasma (**f**), or CM from A4573 cells (**g**) and other indicated EwS cell lines (**h**), using the indicated Agilent RNA chips. The major bands in cellular RNA preparations correspond to 18S and 28S ribosomal RNAs and tRNAs.

**Extended Data Figure 2.**
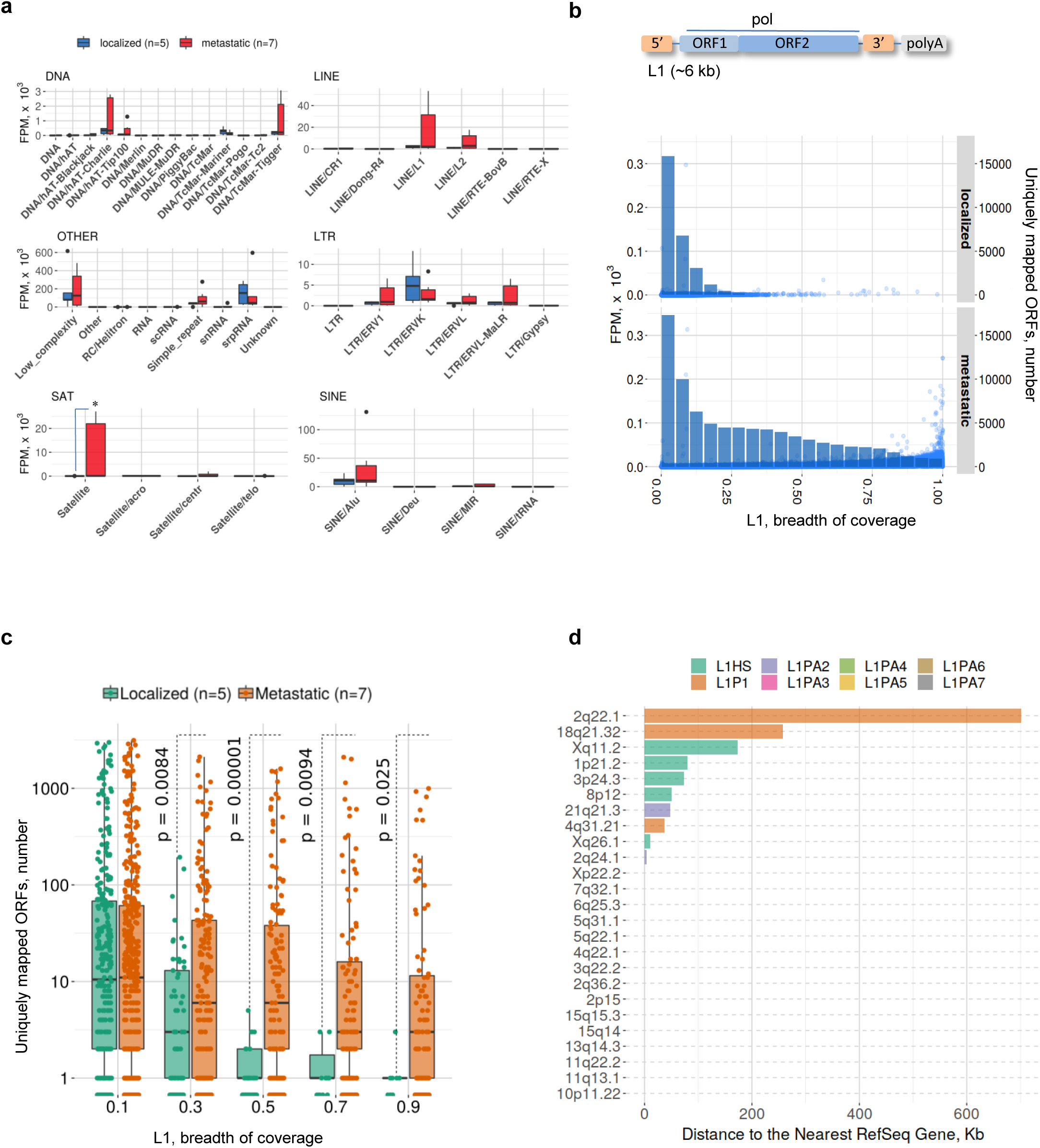
Representation of repeat RNAs in plasma EVs of EwS patients, *L1* protein-coding potential and genomic locations. **a**, Representation of RepeatMasker-annotated RNAs in plasma EVs from EwS patients with localized (n=5) or metastatic disease (n=7). Data are mean FPM ± SEM; *p<0.05, one-tailed unpaired t-test. **b,** Outline of *L1* structural organization (top) and scatter plots of RNAseq read coverage (bottom) of *L1* ORFs (annotated as “pol”; gEVE database http://geve.med.u-tokai.ac.jp). The Y-axis, number of uniquely mapped ORFs; the X-axis, breadth of coverage (length covered/total length of the entire element). Histograms, number of unique L1 ORFs with different breadths of coverage. **c,** Box plots, number of unique *L1* ORFs at different breadth of coverage from (**b**). Comparisons made between the number of ORFs in metastatic vs localized; p-values were calculated by Wilcoxon rank sum test. **d,** Genomic locations and distance to the nearest gene for the top 25 *L1* RNAs from (**b**) ORFs are predicted to encode polypeptides >300 aa. Data in (**b**), (**c**) and (**d**) are linked to Extended Data Table 5a.

**Extended Data Figure 3.**
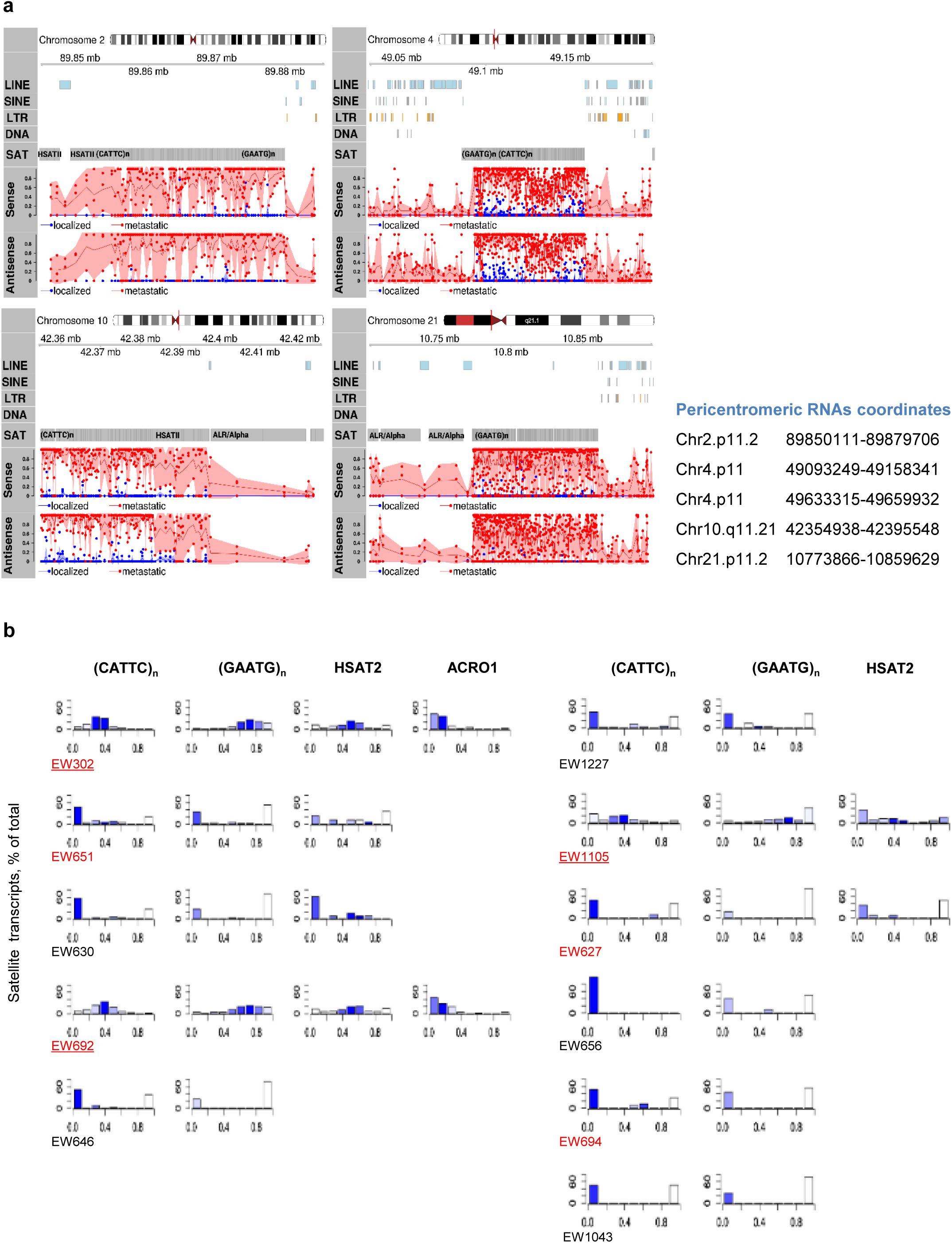
Plasma EVs from EwS patients with metastasis are enriched with sense and antisense pericentromeric RNAs from defined genomic loci. **a**, Coverage of four major genomic loci with sense and antisense satellite RNAseq reads. EwS patients with metastasis or localized tumors are marked red and blue, respectively. **b,** Strand-specific alignment of RNAseq reads shown in (**a**). The Y-axis, percentage of transcripts in sense or antisense orientation; the X-axis, a Jaccard coefficient calculated as antisense/total reads. Values close to 0.5 indicate equal proportion of sense and antisense transcripts, while ratios close to zero or 1 represent transcripts in sense or antisense orientation, respectively. Read abundance in each bin is reflected by color coding, with dark blue indicating higher coverage. No bar plots are shown for cases with no reads.

**Extended Data Figure 4.**
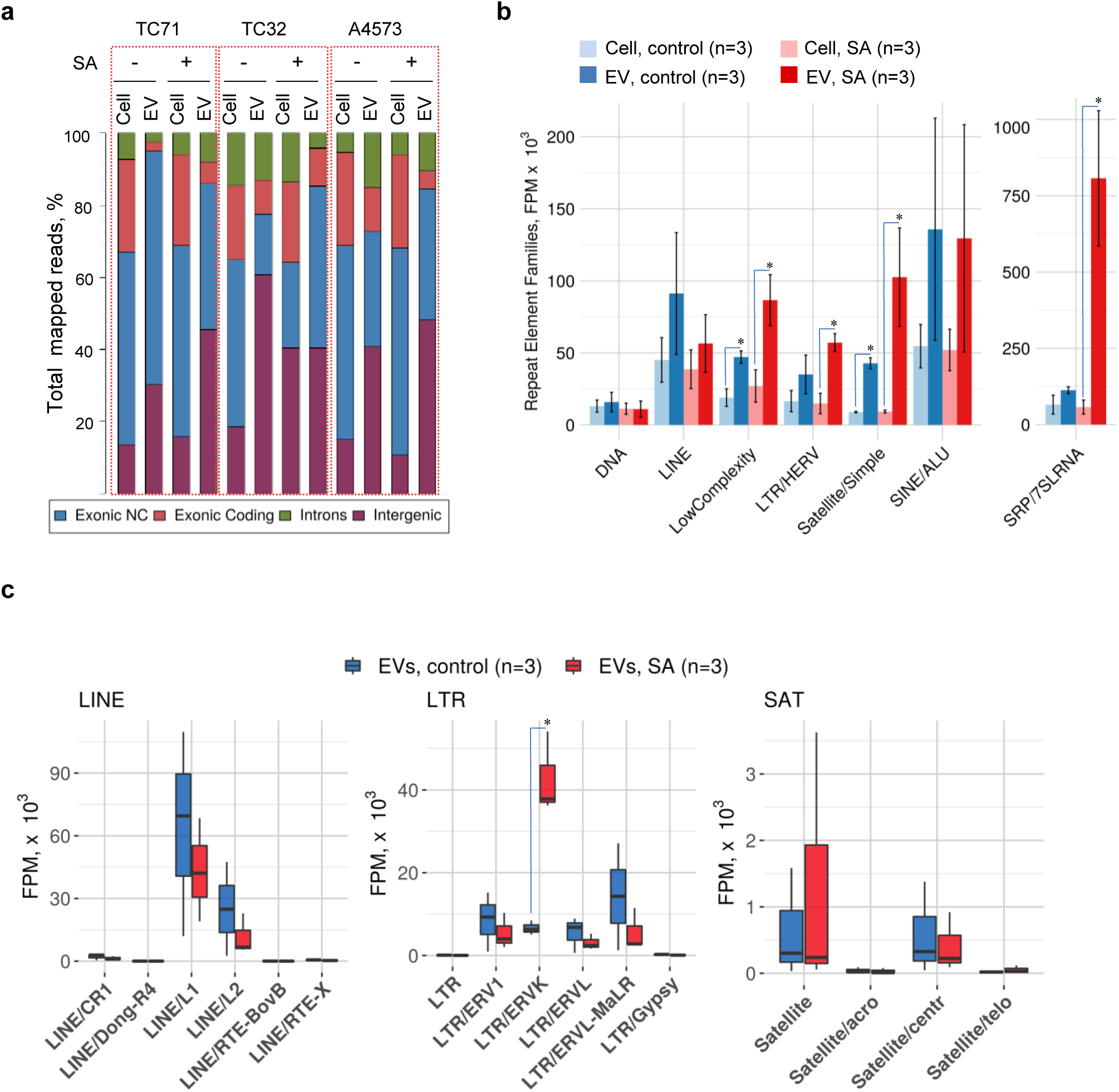
EVs from EwS cell lines are enriched in repeat RNAs and depleted of mRNAs. **a**, Distribution of normalized non-ribosomal RNAseq reads identified in three indicated EwS cell lines treated with vehicle control (-) or sodium arsenate (SA). Exonic non-coding (NC), exonic coding, intronic and intergenic transcripts are shown. **b,** Representation of repeat RNA families in combined cell line (Cell) and EV datasets from (**a**). Data are mean ± SEM (n=3); *p<0.05, Wilcoxon rank sum test. Data are linked to Extended Data Table 4. **c,** Representation of RepeatMasker-annotated RNAs in EVs from EwS cell lines treated with PBS (control) or SA from (**b**); *p<0.05, one-tailed unpaired t-test.

**Extended Data Figure 5.**
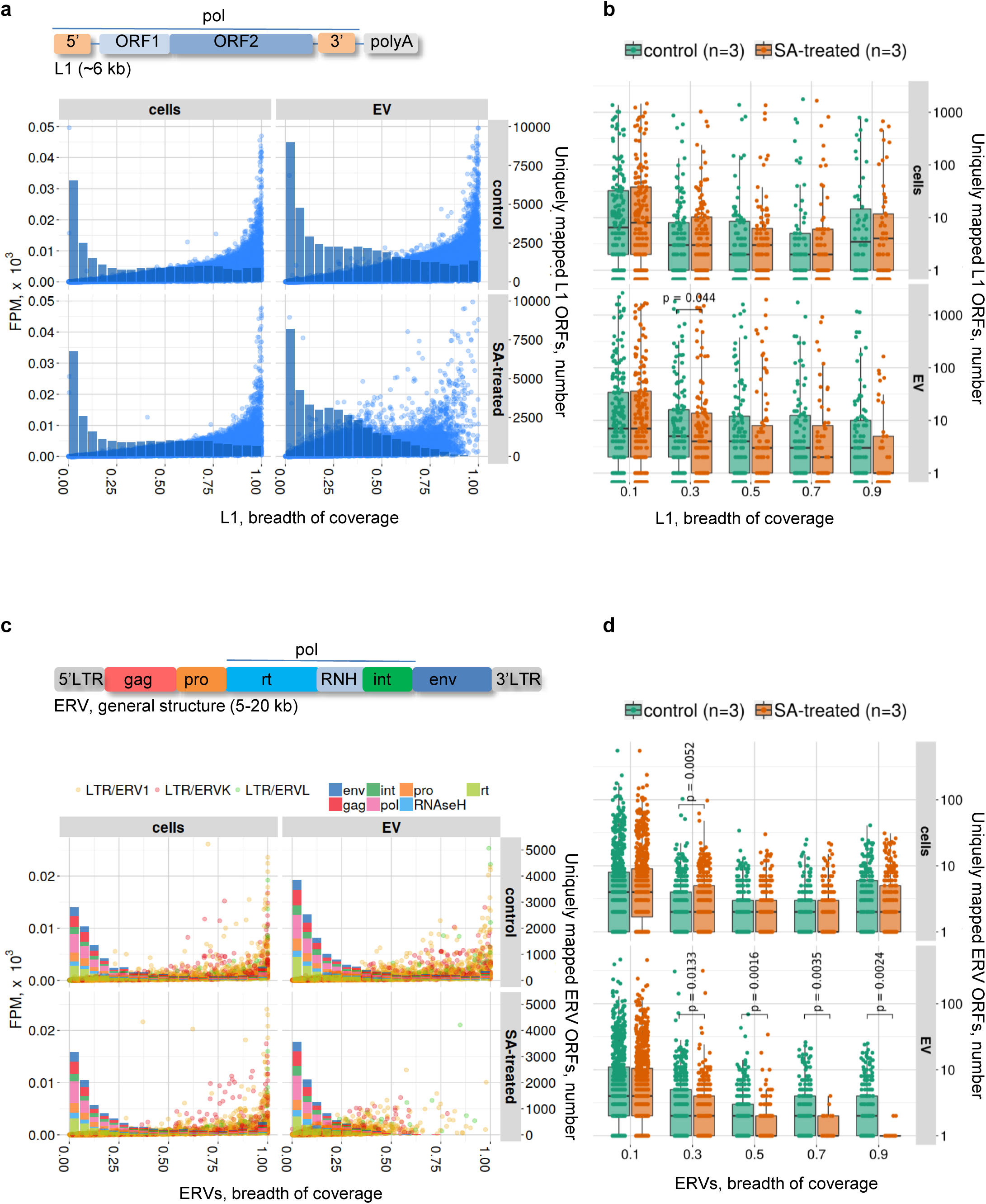
RNAseq read coverage of *L1* and *ERV* ORFs. **a, c,** Scatter plots of RNAseq read coverage of *L1* (**a**) and indicated *ERV* (**c**) ORFs detected in A4573, TC32 and TC71 cells and their EVs; combined datasets for cells and EVs are shown. The Y-axis, number of uniquely mapped ORFs; the X-axis, breadth of coverage (length covered/total length of the locus). Histograms, number of unique ORFs at different breadths of coverage. **b, d**, Box plots, number of unique ORFs with different breadth of coverage from (**a, c**). Comparisons made with mock-treated cells; p-values calculated by Wilcoxon rank sum test. Data are linked to Extended Data Table 5b.

**Extended Data Figure 6.**
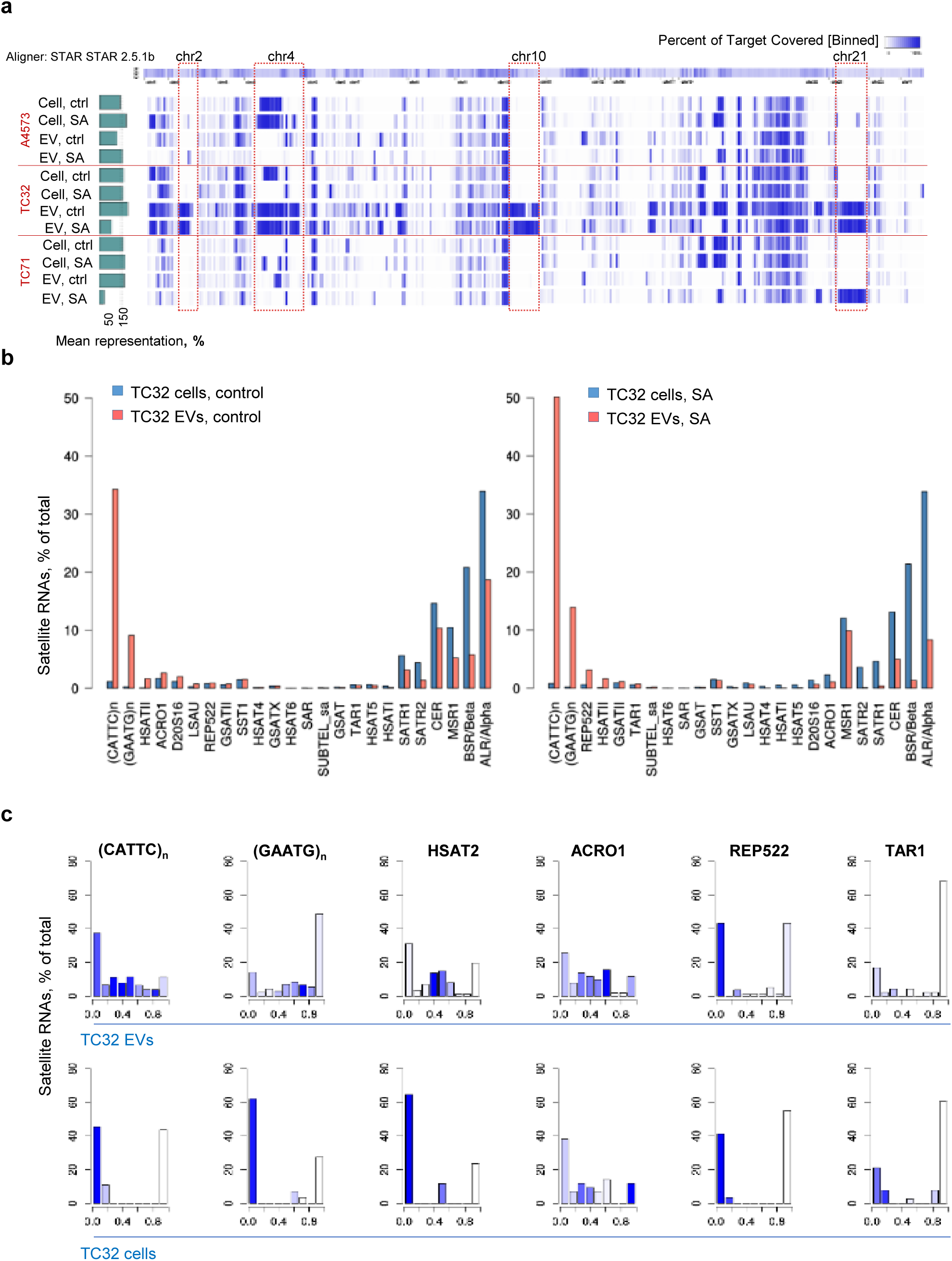
TC32 EVs are enriched with chr. 2, 4, 10 and 21 pericentromeric transcripts. **a**, Genome-wide mapping of satellite RNAseq reads derived from EwS cell lines treated with SA or vehicle control (ctrl) or their respective EVs, as indicated. Blue color intensity shows breadth of coverage. Green bars to the left indicate total read counts for each sample expressed as a percentage of mean. **b,** Representation of various satellite RNAs detected in TC32 cells, vehicle control or SA-treated, and their respective EVs. Proportion of each family is expressed as a percentage of total satellite reads. **c,** Strand-specific alignment of satellite reads derived from mock-treated TC32 cells and their EVs. The Y-axis, percentage of satellite transcripts in sense or antisense orientation; the X-axis, a Jaccard coefficient calculated as antisense/total reads. Values close to 0.5 indicate equal proportion of sense and antisense transcripts, supporting a presence of complementary sequences, while ratios close to zero or 1 represent transcripts in sense or antisense orientation, respectively. Read abundance in each bin is reflected by color, with dark blue indicating higher coverage.

**Extended Data Figure 7.**
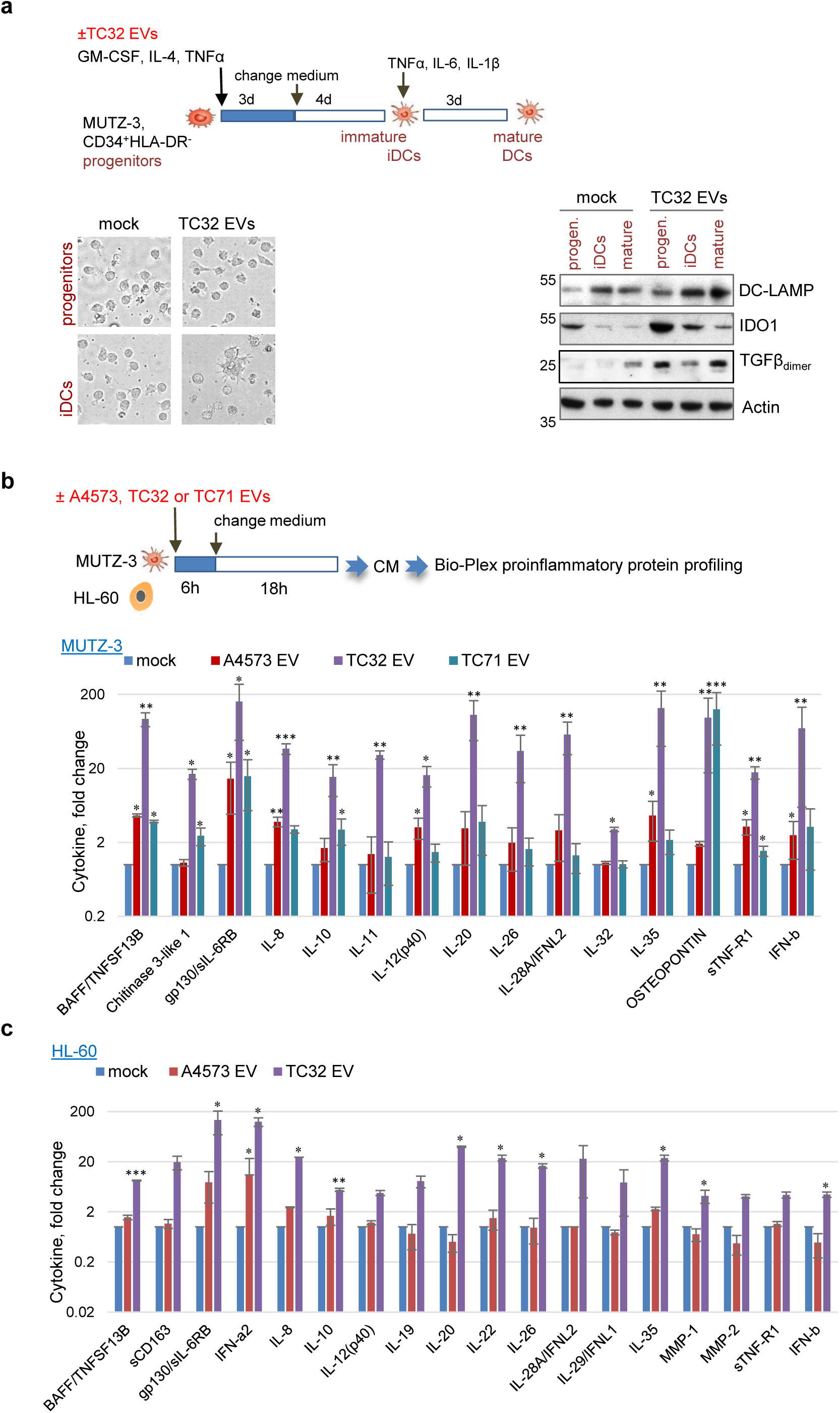
EwS EVs alter differentiation and induce proinflammatory responses in myeloid progenitor cells. **a**, Outline of MUTZ-3 differentiation experiments (top), phase contrast cell imaging and immunoblotting analysis (bottom) for the indicated proteins (representative of three independent experiments). Undifferentiated progenitor cells (progen.), immature (iDCs) and mature DCs are shown. **b**, **c**, Outline of MUTZ-3 and HL-60 cell treatment experiments and Bio-Plex 37 proinflammatory protein profiling of the CM from MUTZ-3 (**b**) or HL-60 (**c**) cells treated with EVs from the indicated cells or mock. Proteins detected in CM from EV- vs mock-treated cells (set as 1 for each cytokine) above a 2-fold threshold in at least one treatment condition are shown. Data are mean ± SD (n=3); *p<0.05, **p≤0.01, ***p<0.001, paired 2-tailed t-test.

**Extended Data Figure 8.**
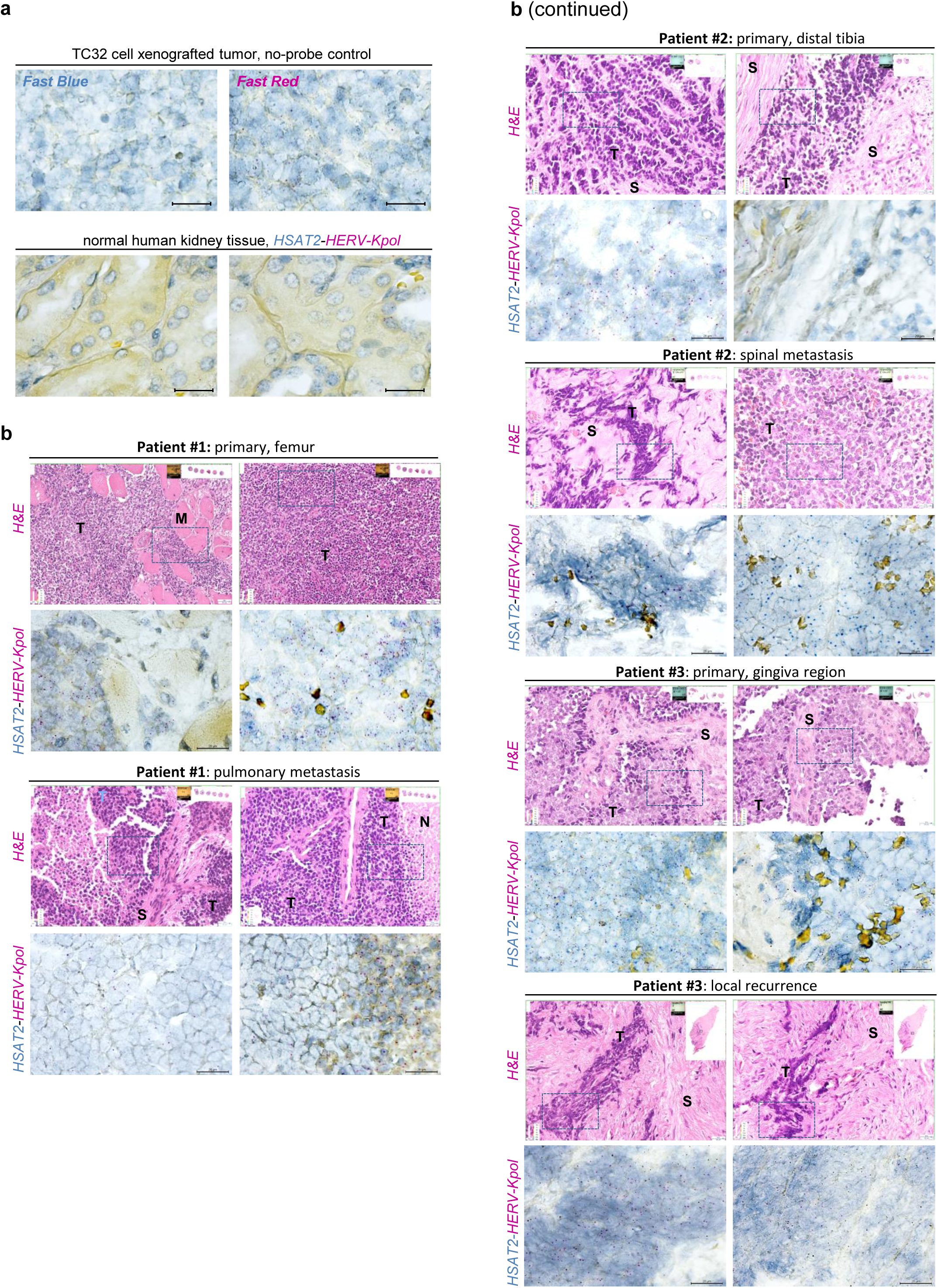
Representative images of *HSAT2* and *HERV-Kpol* RNA detected in matching primary and metastatic tumors from three EwS patients. **a**, ViewRNA ISH staining to confirm signal specificity. Top, no-probe control; staining of TC32 xenografted tumor section was performed in the absence of probes and developed with Fast Blue or Fast Red substrates, as indicated. Bottom, normal human kidney tissue staining with the HSAT2 and HERV-Kpol probes. **b**, Representative images of H&Es (top panels, x40 magnification) and complementary ViewRNA ISH staining (bottom panels, x100 magnification) of primary and metastatic (or recurrent) tumors from the same patient (n=3). ViewRNA images were taken from areas denoted by squares. The inserts on each H&E image show the slide layout and position of the image captured. T- tumor; M- muscle cells; S- stroma; N- necrosis. Scale bars, 20 μm.

**Extended Data Figure 9.**
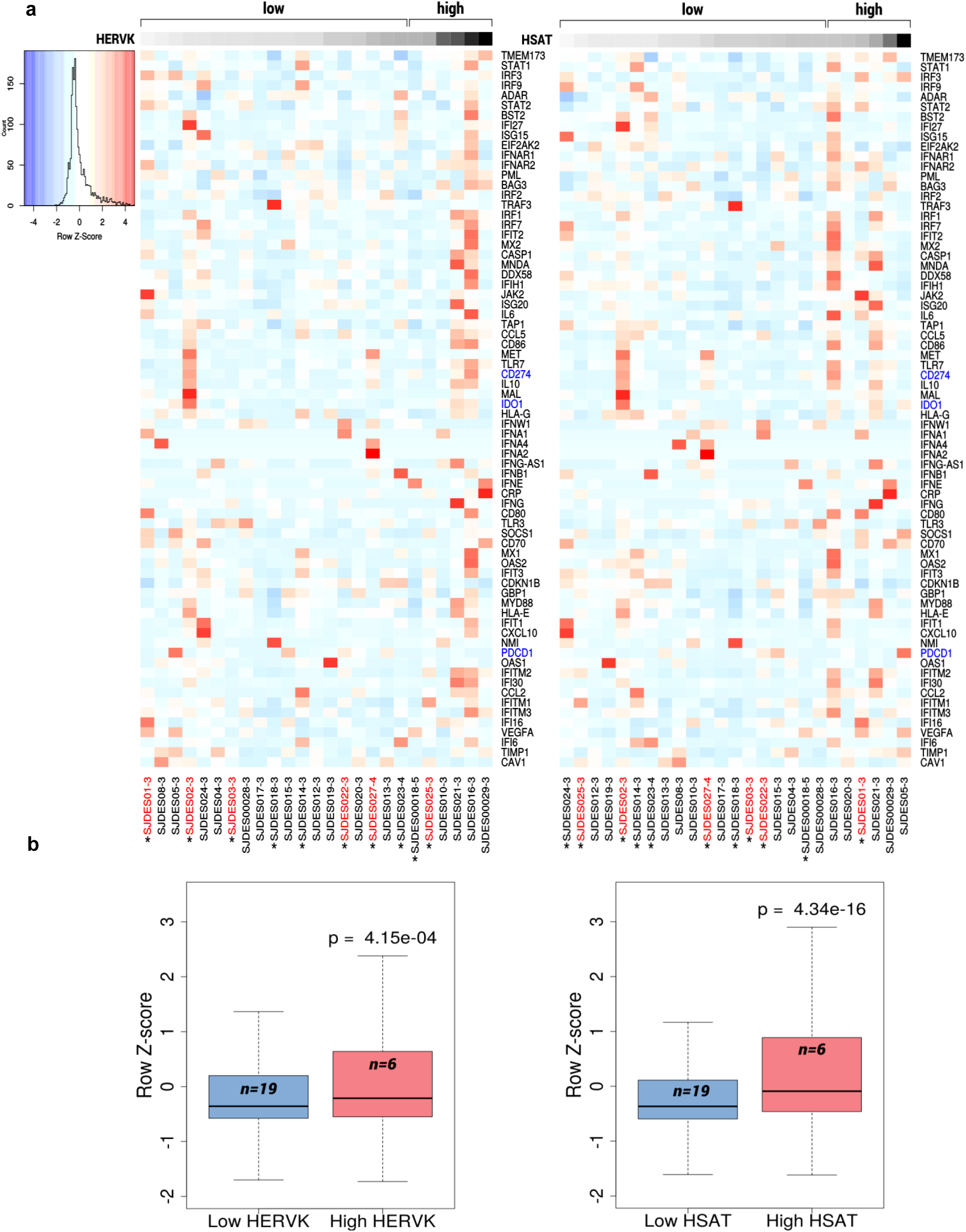
*HERV-K* and *HSAT* RNAs expression is strongly associated with increased expression of IFN-stimulated viral response transcripts. **a,** Heatmap of the IFN-stimulated gene expression and *CD274/PD-L1*, *PDCD1/PD-1* and *IDO1* (shown in blue) in 25 EwS tumors from the dbGaP RNAseq dataset generated by Crompton et al., 2014. Increasing levels of *HERV-K* or *HSAT* (represented by combined *HSAT2*, *(CATTC)_n_* and *(GAATG)_n_* reads) in each sample are shown on the top as gray scale gradient. Samples are split into “low” (n=19) or “high” (n=6) groups based on their HERV-K/HSAT coverage relative to the mean value across all samples. Patients with metastasis are marked red. Star denotes post-chemotherapy patients. **b**, Box-and-whisker plots of IFN-inducible transcript expression values in “low” and “high” HERV-K/HSAT expression groups from (**a**); p-values were calculated using Wilcoxon rank sum test.

**Extended Data Figure 10.**
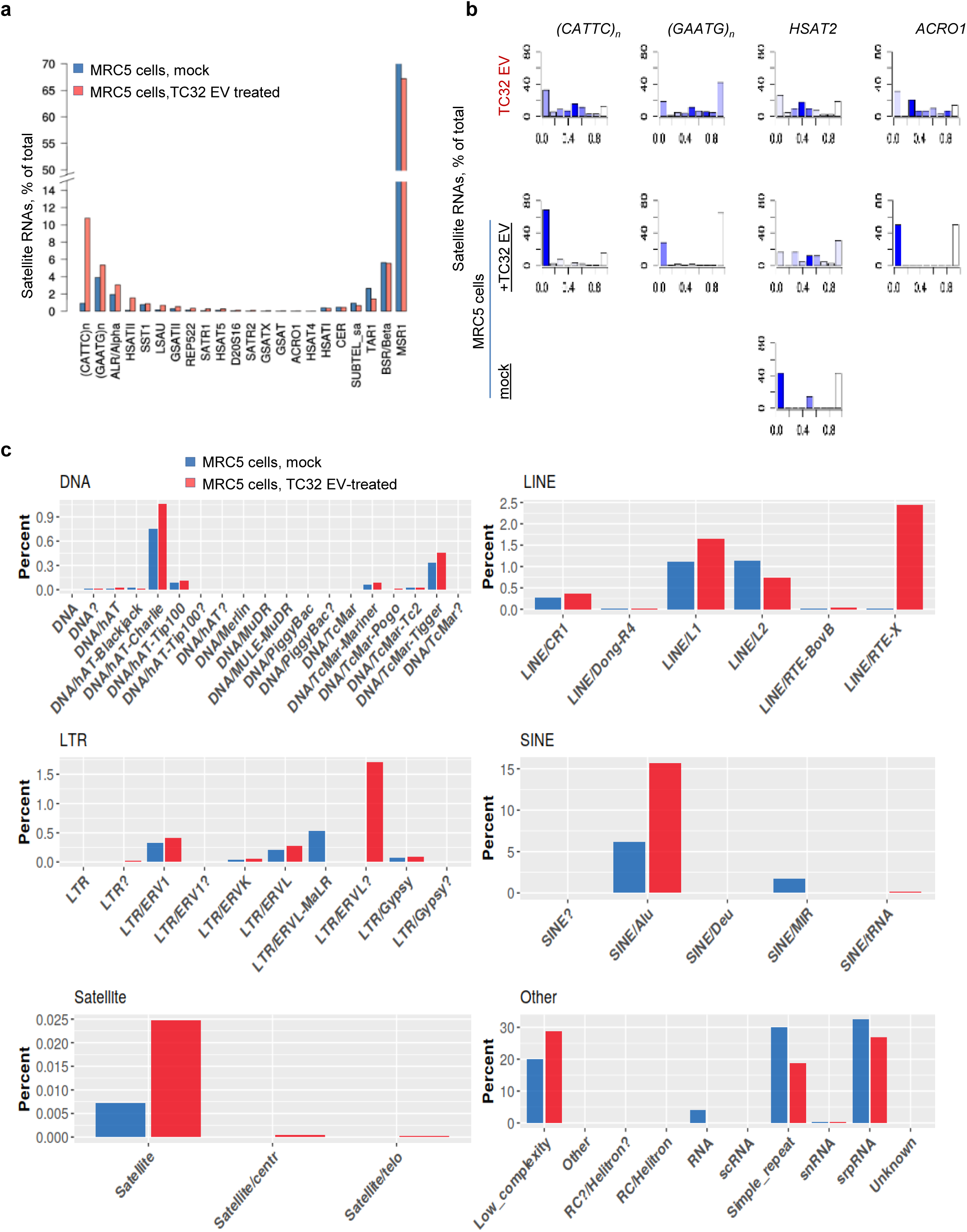
Treatment with TC32 EVs induces selective accumulation of repeat RNA subsets in the recipient MRC5 cells. **a**, Representation of RepeatMasker-annotated RNAs from various satellite families in MRC5 cells treated with mock or TC32 EVs. **b,** Strand-specific alignment of satellite RNAseq reads derived from mock- or TC32 EV-treated MRC5 cells; TC32 EVs used for treatment are marked red. The Y-axis shows percentage of transcripts from the satellite repeat families; the X-axis, Jaccard coefficient calculated as antisense/total reads; values close to 0.5 indicate equal proportion of sense and antisense transcripts, while ratios close to zero or 1 represent transcripts in sense or antisense orientation, respectively. Read abundance in each bin is reflected by color coding, with dark blue indicating higher coverage. No bar plots are shown for cases with no reads. **c,** Representation of RepeatMasker-annotated RNAs in mock- or TC32 EV-treated MRC5 cells.

**Extended Data Figure 11.**
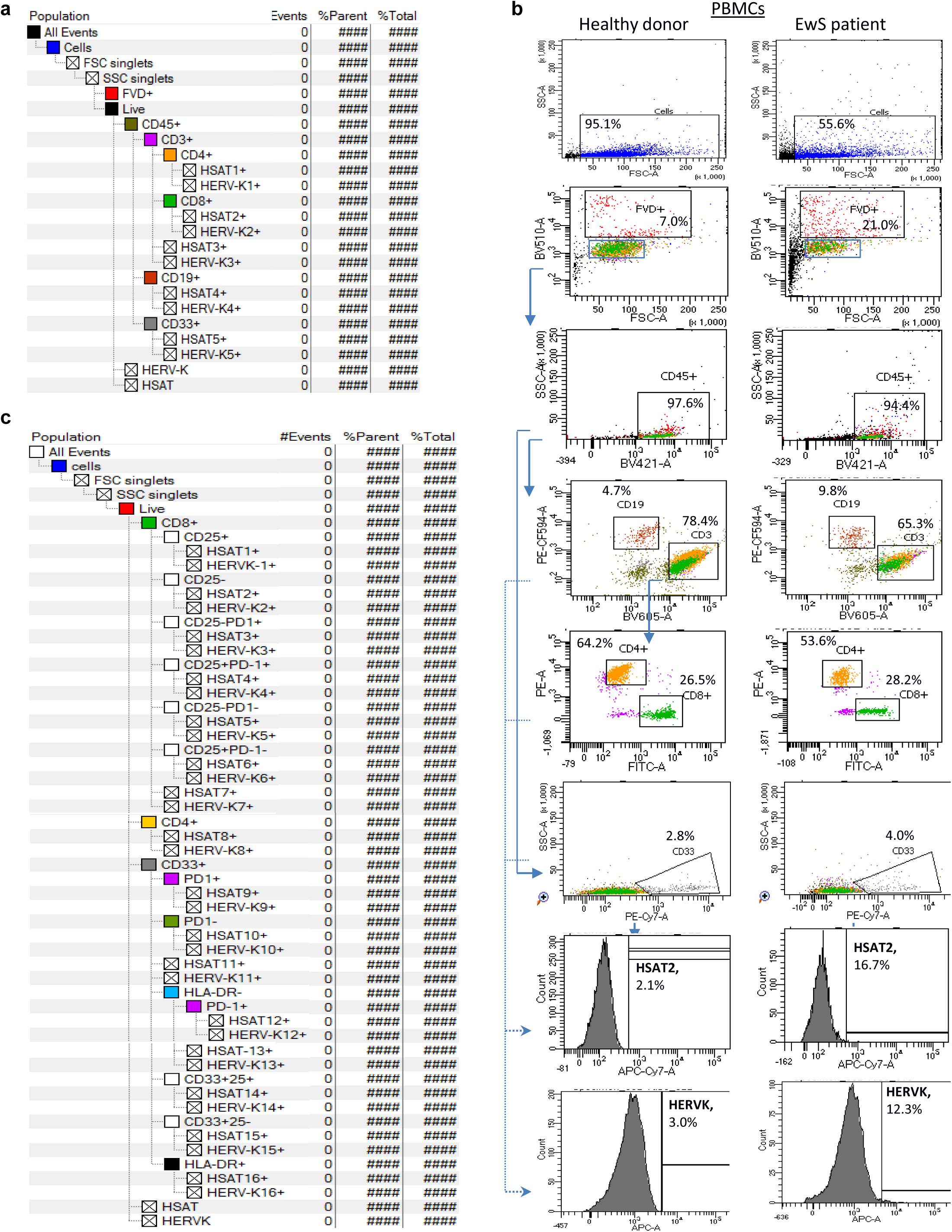
PBMC PrimeFlow RNA assay and gating strategies. **a, b**, Gating strategy template (**a**) for the results shown in Fig. 3 a, c, and representative results obtained from healthy donor and EwS patient PBMCs (**b**). **c,** Gating strategy template for the results shown in Fig. 3b. Doublets were excluded and live FVD-negative cells (denoted by a blue square in (**b**)) were analyzed, as indicated. Signal specificity was defined by the respective FMO controls and isotope control antibodies. Percentage of *HERV-K* and *HSAT2*-positive cells was measured in each cell population.

## SUPLEMENTARY TABLES

**Extended Data Table 1. List of EwS patients, their clinical characteristics and plasma levels of indicated cytokines and transcripts**

**Extended Data Table 2. List of healthy donors and plasma levels of indicated cytokines and transcripts**

**Extended Data Table 3a. Multivariate analysis showing association between plasma cytokine levels and clinical parameters**

**Extended Data Table 3b. Multivariate analysis showing association between the indicated plasma EV transcripts and clinical parameters**

**Extended Data Table 4. Summary of RNAseq data**

**Extended Data Table 5a. List of ERV and L1 RNAs with predicted long ORFs (encoding polypeptides >80 amino acids) identified in plasma EVs of EwS patients (linked to Fig. 2d, e and Extended Data Fig. 2b-d)**

**Extended Data Table 5b. List of ERV and L1 RNAs with predicted long ORFs (encoding polypeptides >80 amino acids) identified in A4573, TC32 and TC71 EwS cell lines and the respective EVs (linked to Extended Data Fig. 4a,b)**

**Extended Data Table 6a. List of 109 mRNAs downregulated in TC32 EV-treated MRC5 cells compared to mock and their GO annotations**

**Extended Data Table 6b. List of 297 mRNAs upregulated in TC32 EV-treated MRC5 cells compared to mock and their GO annotations**

**Extended Data Table 6c. Reactome Functional Interaction (FI) network analysis of 54 mRNAs from “Chromatin-remodeling”, “Cellular response to DNA damage” and “Cell division” GO categories upregulated in TC32 EV-treated MRC5 cells compared to mock**

**Extended Data Table 7. List of antibodies and reagents used for immunoblotting, PrimeFlow RNA and ViewRNA assays**

**Extended Data Table 8. Cell lines, their characteristics and growth conditions Extended Data Table 9. RT-PCR primers used in this study**

